# Myosin 10 uses its MyTH4 and FERM domains differentially to support two aspects of spindle pole biology required for mitotic spindle bipolarity

**DOI:** 10.1101/2023.06.15.545002

**Authors:** Yang-In Yim, Antonio Pedrosa, Xufeng Wu, Krishna Chinthalapudi, Richard E. Cheney, John A. Hammer

**Affiliations:** Cell and Developmental Biology Center, National Heart, Lung and Blood Institute, National Institutes of Health, Bethesda, MD; Department of Physiology and Cell Biology, College of Medicine, The Ohio State University, Columbus, OH; Department of Cell Biology and Physiology, University of North Carolina, Chapel Hill, NC

## Abstract

Myosin 10 (Myo10) has the ability to link actin filaments to integrin-based adhesions and to microtubules by virtue of its integrin-binding FERM domain and microtubule-binding MyTH4 domain, respectively. Here we used Myo10 knockout cells to define Myo10’s contribution to the maintenance of spindle bipolarity, and complementation to quantitate the relative contributions of its MyTH4 and FERM domains. Myo10 knockout HeLa cells and mouse embryo fibroblasts (MEFs) both exhibit a pronounced increase in the frequency of multipolar spindles. Staining of unsynchronized metaphase cells showed that the primary driver of spindle multipolarity in knockout MEFs and knockout HeLa cells lacking supernumerary centrosomes is pericentriolar material (PCM) fragmentation, which creates y-tubulin-positive acentriolar foci that serve as additional spindle poles. For HeLa cells possessing supernumerary centrosomes, Myo10 depletion further accentuates spindle multipolarity by impairing the clustering of the extra spindle poles. Complementation experiments show that Myo10 must interact with both integrins and microtubules to promote PCM/pole integrity. Conversely, Myo10’s ability to promote the clustering of supernumerary centrosomes only requires that it interact with integrins. Importantly, images of Halo-Myo10 knock-in cells show that the myosin localizes exclusively within adhesive retraction fibers during mitosis. Based on these and other results, we conclude that Myo10 promotes PCM/pole integrity at a distance, and that it facilitates supernumerary centrosome clustering by promoting retraction fiber-based cell adhesion, which likely provides an anchor for the microtubule-based forces driving pole focusing.

## Introduction

Myosin 10 (Myo10) is a member of the MyTH4/FERM domain family of unconventional myosins (reviewed in [1-5]). These actin-based motors are special in that they can interact with microtubules and integrins via their MyTH4 and FERM domains, respectively. Myo10 is known as the “filopodial myosin” because it accumulates dramatically at the tips of filopodia [6-8], and because cells that over-express or lack Myo10 exhibit increases and decreases in filopodia number, respectively [6, 9-18]. Consistent with these observations, Myo10 prefers to walk on bundled actin filaments like those that comprise filopodia [19-21], and it can be seen to move out filopodia at ∼800 nm/s [6, 22]. Perhaps most importantly for this study, Myo10 promotes the formation of adhesions within filopodia by virtue of its FERM domain-dependent binding to β1-integrin [23-25]. These filopodial adhesions can sense extracellular matrix (ECM) stiffness and topography [26-29], generate traction force [9, 26, 28, 30-33], and often mature into focal adhesions upon cell advance [27, 29, 34-38]. Consistently, Myo10-dependent filopodial adhesions have been implicated in cell migration and cancer cell metastasis (reviewed in [9, 39, 40]).

While the bulk of cell biological studies to date have focused on Myo10’s role in the structure and function of filopodia, which are generally devoid of microtubules, Myo10 also plays important roles in microtubule-dependent processes. The best evidence for this has come from studies focusing on Myo10’s role in meiosis and mitosis. First, Bement and colleagues showed that Myo10 localizes to the portion of the frog egg meiotic spindle in contact with the actin cortex, and that disrupting Myo10 function inhibits nuclear anchoring, spindle assembly and spindle-cortex association in eggs [41]. Subsequently, Bement and colleagues reported that Myo10 localizes to spindles and spindle poles in embryonic frog epithelial cells undergoing mitosis, and that morpholino-based partial knockdown of Myo10 causes defects in spindle anchoring, spindle length, spindle dynamics, and metaphase progression [42]. They also showed that about 15% of spindle poles in Myo10 morphants fragment during early anaphase, leading to multipolar spindles [42]. While the extent to which this apparent defect in pole stability was due to centriole disengagement versus PCM fragmentation was not determined, and while a requirement for MyTH4 domain: microtubule interaction was not demonstrated for any of the phenotypes, Bement and colleagues did show that Myo10’s isolated MyTH4/FERM domain interacts with the pole factor TPX2, and that Myo10 is required for the robust localization of TPX2 at and near poles. Based on these findings, they argued that Myo10 maintains spindle pole integrity by recruiting TPX2, and that, without the stabilizing effect of TPX2, poles in Myo10 knockdown cells tend to fragment when subjected to the strong chromosomal and spindle forces that ramp up during metaphase.

Recent work has shown that Myo10 also plays a role in positioning the mitotic spindle [43]. In this study, Pellman and colleagues presented evidence that Myo10 localizes to subcortical actin clouds at the equator of dividing HeLa cells where it cooperates in a non-redundant fashion with cortical dynein to position the mitotic spindle. Importantly, Myo10 must be capable of interacting with microtubules to assist in spindle positioning, as the positioning defect exhibited by Myo10 knockdown cells was not rescued by a version of Myo10 harboring a mutated MyTH4 domain that can no longer bind to microtubules. That said, Toyoshima and colleagues [44] showed in an earlier study that the ability of integrin-based adhesions to orient the mitotic spindle parallel to the substratum requires Myo10. That result suggests Myo10 might promote spindle positioning at least in part through its FERM domain-dependent interaction with integrins, although this possibility was not explored. In a subsequent study, however, Toyoshima and colleagues [45] presented evidence the that cyclin-dependent kinase PCTK1 phosphorylates KAP0, a regulatory subunit of protein kinase A, that phospho-KAP0 binds to the FERM domain of Myo10 to enhance its interaction with integrin, and that this enhancement facilitates the retraction fiber-dependent control of spindle orientation [46, 47].

Finally, Myo10 has been implicated along with the pole-focusing, microtubule minus end-directed kinesin 14 family member HSET in the clustering of the extra spindle poles that appear in cells exhibiting supernumerary centrosomes during interphase [48]. This process, which is usually referred to as supernumerary centrosome clustering, serves to cluster the extra spindle poles into two groups to enable a bipolar mitosis. Myo10 may cooperate with HSET to promote the clustering of supernumerary centrosomes in a manner similar to how it cooperates with cortical dynein to position the mitotic spindle [43], as Myo10 with a mutated MyTH4 domain could not rescue the multipolar phenotype exhibited by Myo10 knockdown cells over overexpressing PLK4 [43]. Whether Myo10 must also interact with integrins via its FERM domain to promote supernumerary centrosome clustering has not been tested.

Here we used Myo10 knockout (KO) HeLa cells created using CRISPR and MEFs isolated from our Myo10 KO mouse [13] to define the contribution that Myo10 makes to maintaining spindle bipolarity. We then used complementation of HeLa KO cells with mutated versions of Myo10 to quantitate the contributions that its microtubule-binding MyTH4 domain and integrin-binding FERM domain make to the myosin’s ability to maintain spindle bipolarity. We find that both cell types exhibit a pronounced increase in the frequency of multipolar spindles, and that two separate defects related to spindle pole biology are responsible for this when Myo10 is missing. The first defect, spindle pole fragmentation, is the major driver of multipolar spindles in knockout MEFs and knockout HeLa cells lacking supernumerary centrosomes. We show that pole fragmentation is due almost entirely to PCM fragmentation (i.e. centriole disengagement plays only a minor role), and that fragmentation occurs as cells approach metaphase, arguing that it is a force-dependent event. We do not, however, see an obvious defect in pole maturation, as the pole localization of both TPX2 and a standard marker for pole maturation are normal in the absence of Myo10. Moreover, we do not see Myo10 at poles in cells where one Myo10 allele was tagged with Halo using CRISPR. Together, these results indicate that the defect in PCM integrity is not part of a general defect in pole maturation, and they suggest that Myo10 promotes PCM/pole integrity at a distance (for example, through its regulation of the CDK1 kinase inhibitor WEE1 [49]). The second defect, an inability to cluster extra spindle poles, is the major driver of multipolar spindles in knockout HeLa cells possessing supernumerary centrosomes. Unlike the defect in PCM/pole stability, where complementation shows that the Myo10 must interact with both integrins and microtubules to promote stability, Myo10 only needs interact with integrins to promote supernumerary centrosome clustering. Consistently, endogenously tagged Myo10 localizes dramatically within mitotic cells to retraction fibers, which are known to be required for the integrin-dependent adhesion of mitotic cells [50-55]. These and other results argue that Myo10 promotes the clustering of supernumerary centrosomes by promoting retraction fiber-based cell adhesion during mitosis, which likely serves as one of several anchors required for the efficient focusing of extra poles by microtubule-based forces [56]. Given that supernumerary centrosomes are rare in normal cells but common in transformed cells, and that failure to cluster the ensuing extra spindle poles results in a multipolar division, aneuploidy, and cell death [57-59], inhibiting Myo10’s contribution to this process should selectively kill cancer cells.

## Results

### Myo10 KO HeLa cells exhibit a pronounced increase in the frequency of multipolar spindles

We used CRISPR-Cas9-based genome editing in an effort to abrogate Myo10 expression in HeLa cells, as these cells have been used extensively to explore Myo10 functions, including those related to mitosis. Of note, HeLa cells are polyploid (they contain 70-90 chromosomes instead of the 46 in normal human cells), and they appear to contain at least three Myo10 alleles based on analyses of chromosomal duplications [60]. We inserted guide sequences targeting Exon 3 of the Myo10 gene into plasmid pSpCas9(BB)-2A-GFP to create the construct pSpCas9(BB)-2A-GFP-ghM10-Exon3, which we then introduced into HeLa cells by electroporation. Two days later, GFP-positive cells were subjected to single-cell sorting into 96- well plates. This effort yielded only two putative Myo10 KO clones (KO-1 and KO-2) from 384 wells, or ∼0.5% of sorted cells. This low number likely reflects several factors, including the stresses incurred by single-cell sorting (only ∼30% of control cells sorted to single cells grew up), the defects in mitosis that occur when Myo10 is depleted (see below), and the fact that these defects are more pronounced at lower cell densities (see below). Western blots of whole cell extracts prepared from WT HeLa, KO-1 and KO-2 and probed with an antibody to Myo10 showed a large reduction in Myo10 levels in KO-1 and KO-2 (Fig. 1A), with scans of replicate blots showing that these cells contain 7.1% and 13.8% of normal Myo10 levels, respectively (Fig. S1A). Sequencing of PCR products that span the portion of Exon 3 targeted by the guide sequences revealed two frameshift alleles and one missense allele in both KO-1 and KO-2. We conclude, therefore, that both KO lines likely contain two nonsense alleles and one missense allele, making them hypomorphs for Myo10. While not complete KOs, we refer to these two lines as KO lines for the sake of simplicity.

**Figure 1.**
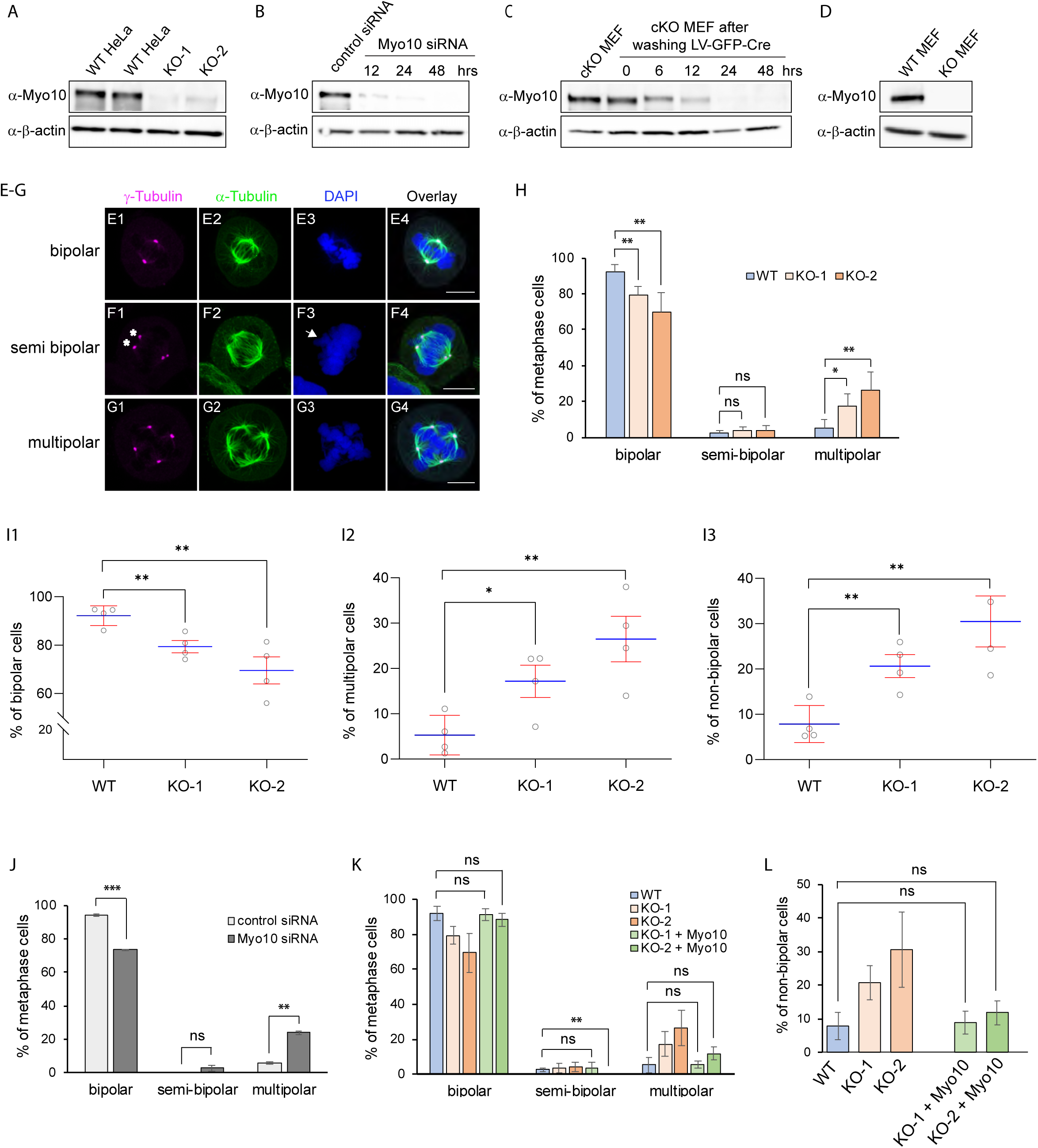
Myo10 depleted cells exhibit a significant increase in the frequency of multipolar spindles at metaphase. (A-D) Westerns blots of whole cell extracts prepared from WT and Myo10 KO HeLa cells (A), HeLa cells treated twice for 48 hrs each with a control, nontargeting siRNA or twice with a Myo10 Smartpool siRNA and collected 12, 24 and 48 hrs after starting the second treatment (B), non-transduced cKO MEFs and cKO MEFs 0, 6, 12, 24 and 48 hrs after transduction with lentiviral cre (C), and MEFs from a WT B6 mouse and from the tm1d Myo10 KO mouse (D) probed for Myo10 and β-actin as a loading control. (E1-E4) A representative example of a metaphase HeLa cell harboring a bipolar spindle stained for γ-tubulin, α-tubulin, and DNA (DAPI), along with the overlay (see also Z-Stack Movie 1). (F1-F4) Same as E1-E4 except that this HeLa cell harbored a semi-bipolar spindle (see also Z-Stack Movie 2). Two γ- tubulin-positive poles with asterisk indicate they were in different z planes, resulting in chromosome protrusion (arrow in F3). (G1-G4) Same as E1-E4 except that this HeLa cell harbored a multipolar spindle (see also Z-Stack Movie 3). (H) Quantitation at metaphase of the percent of bipolar, semi-bipolar, and multipolar spindles in WT HeLa (281 cells) and HeLa Myo10 KO lines KO-1 (289 cells) and KO-2 (307 cells) from 4 experiments. (I1-I3) Comparison of WT Hela with KO-1 and KO-2 with regard to percent bipolar cells (I1), percent multipolar cells (I2) and percent non-bipolar cells (semi-bipolar plus multipolar), all at metaphase (calculated from the raw data in (H); shown as SEMs). (J) Quantitation at metaphase of the percent of bipolar, semi-bipolar, and multipolar spindles in HeLa cells treated twice (each for 48 hrs) with a control, nontargeting siRNA or with Myo10 Smartpool siRNA (173 control cells and 234 KD cells from 2 experiments). (K) Quantitation at metaphase of the percent of bipolar, semi-bipolar, and multipolar spindles in WT HeLa, KO-1, and KO-2 cells, and in KO-1 and KO-2 cells 24 hrs post-transfection with mScarlet-Myo10 (103 KO-1 cells and 120 KO-2 cells from 4 experiments; only cells expressing mScarlet-Myo10 were scored; only statistical values for WT versus rescued KO- 1 and KO-2 cells are shown; the values for WT HeLa, KO-1 and KO-2 are from (H)). (L) Quantitation at metaphase of the percent of non-bipolar spindles (semi-bipolar plus multipolar) in WT HeLa, KO-1, and KO-2 cells, and in KO-1 and KO-2 cells 24 hrs post-transfection with mScarlet-Myo10 (only statistical values for WT versus rescued KO-1 and KO-2 cells are shown; the values for WT HeLa, KO-1 and KO-2 are from (H)). All the mag bars are 10 µm.

To identify possible defects in mitotic spindle organization upon Myo10 depletion, we fixed and stained unsynchronized WT HeLa, KO-1 and KO-2 for γ-tubulin to label spindle poles, α- tubulin to label spindle microtubules, and DAPI to label chromosomes. Cells identified as being at metaphase were optically sectioned in 0.25 µm intervals, and the sections used to count spindle pole numbers and to determine spindle and chromosome organization in 3D. Using these images, cells were scored as being either bipolar (this category included not only cells with just two spindle poles, one at each end of a normal-looking bipolar spindle, but also pseudo-bipolar cells where one or both poles at each end of a normal looking bipolar spindle were created by the clustering of supernumerary centrosomes; Fig. 1, E1-E4; Z-Stack Movie 1), semi-polar (this category contained cells possessing two major poles and a normal looking spindle, but also a minor pole in a different focal plane that contributed some microtubules to the spindle; Fig. 1, F1-F4; Z-Stack Movie 2), or multipolar (this category contained cells with more than two major spindle poles; Fig. 1, G1-G4; Z-Stack Movie 3). Scoring of >300 cells per line over four independent experiments showed that KO-1 and KO-2 exhibit 11.8% and 22.7% decreases in the frequency of bipolar spindles, respectively (Fig. IH and I1), 3.2-fold and 5.0-fold increases in the frequency of multipolar spindles (Fig. 1H and I2), and 2.6-fold and 3.9-fold increases in the frequency of non-bipolar spindles (semi-polar plus multipolar) (Fig. 1H and I3). We conclude, therefore, that HeLa cells with reduced levels of Myo10 exhibit a pronounced increase in the frequency of multipolar spindles.

### Myo10 KO HeLa cells exhibit an increase in the frequency of spindles that are not parallel to the substratum

For cells dividing in 2D, the balance of forces driving spindle positioning are such that the spindle’s long axis is usually parallel to the substratum, resulting in the two poles being about equidistant from the substratum i.e. in the same Z-plane. Optical sectioning of ∼200 metaphase cells exhibiting just two poles showed that the average distance in Z separating the two spindle poles was significantly greater in both KO lines than in WT HeLa (Fig. 2A; 2D shows the distributions of these distances in 1 µm intervals; see also Z-Stack Movie 4). This phenotype could well reflect a defect in the adhesion of Myo10 KO cells, resulting in an imbalance in the pulling forces driving spindle positioning ([44]; see Discussion). While the mean values for the separation of the two poles in Z was also elevated in both KO lines at anaphase and telophase (Fig. 2B, 2C; Z-Stack Movies 5 and 6), the differences were not quite statistically significant at anaphase and far from significant at telophase. This result indicates that the defect in positioning the metaphase spindle relative to the substratum seen when Myo10 is depleted is largely corrected as mitosis progresses to telophase.

**Figure 2.**
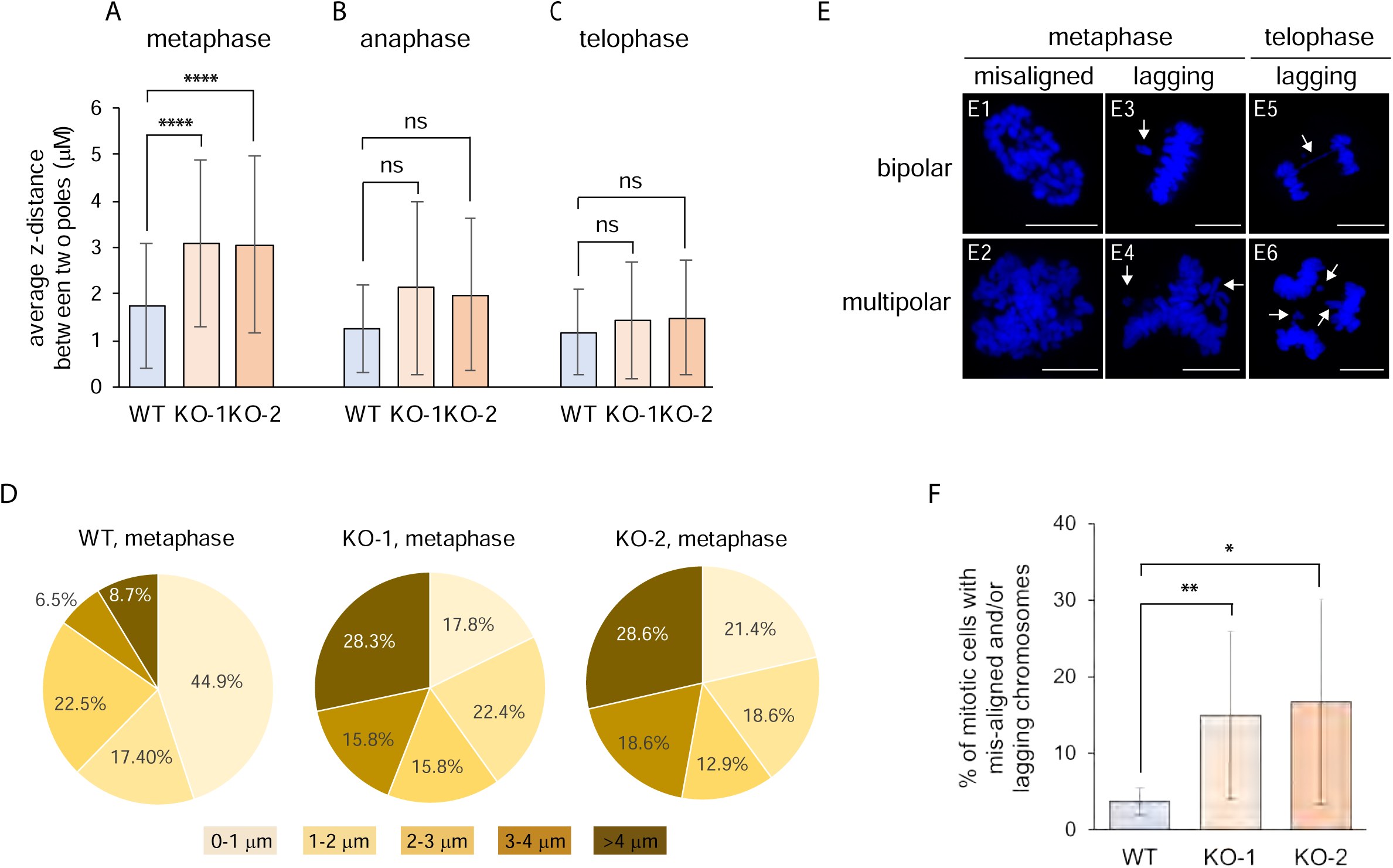
Myo10 KO cells at metaphase exhibit increased frequencies of spindles that are not parallel to the substratum and increased frequencies of misaligned and lagging chromosomes. (A-C) Average distance in Z between the two poles of bipolar WT HeLa, KO-1 and KO-2 at metaphase (A), anaphase (B) and telophase (C) (180 WT cells, 234 KO-1 cells and 228 KO-2 cells from 4 experiments; see also Z-Stack Movies 4-6). (D) The distributions of the values in (A) in 1 um intervals. (E1-E6) Shown are representative examples of misaligned chromosomes at metaphase in KO-1 cells undergoing bipolar mitosis (E1) or multipolar mitosis (E2), and lagging chromosomes at metaphase and telophase in KO-1 cells undergoing bipolar mitosis (E3 and E5, respectively) or multipolar mitosis (E4 and E6, respectively) (arrows point to lagging chromosomes). (F) Percent of mitotic WT HeLa, KO-1 and KO-2 that possess misaligned and/or lagging chromosomes (258 WT cells, 308 KO-1 cells and 330 KO-2 cells from 4 experiments). All mag bars are 10 µm.

### Myo10 KO HeLa cells exhibit increased frequencies of chromosome misalignment and lagging chromosomes

Cells with multipolar or improperly positioned spindles often exhibit chromosome misalignment at metaphase and lagging chromosomes later in mitosis due to merotelic or missing kinetochore attachments. Imaging of mitotic cells following fixation and staining for DNA, microtubules and spindle poles between metaphase and telophase revealed evidence of these chromosome distribution defects in Myo10 KO cells undergoing both bipolar and multipolar divisions (Fig. 2, E1-E6; the arrows in E3-E6 indicate lagging chromosomes). Indeed, scoring showed that the percentage of mitotic cells exhibiting chromosome misalignment and/or lagging chromosomes was 4.0- and 4.5-fold higher in KO-1 and KO-2, respectively, than in WT HeLa cells (Fig. 2F). This result is consistent with the defects in spindle bipolarity and spindle positioning exhibited by these KO lines.

### HeLa cells in which Myo10 levels were reduced using siRNA-mediated knockdown also exhibit an increase in the frequency of multipolar spindles

KO-1 and KO-2 initially grew very slowly, only approaching an approximately normal growth rate about two months in culture (at which point we started collecting data). This behavior suggests that these clonal isolates underwent some kind of genetic or epigenetic change after CRISPR-based genome editing that ameliorated to some extent the cellular defects responsible for their slow growth rate. Given this, and given that both of our KO clones are slight hypomorphs, we decided to also examine HeLa cells in which Myo10 was transiently depleted using siRNA-mediated knockdown (KD). Relative to control cells that received a control, non-targeting siRNA, cells that received two rounds of treatment with Myo10 SmartPool siRNA exhibited near complete depletion of Myo10 48 hours after the second round of treatment (Fig. 1B and Fig. S1B). At 48 hours post-transfection, unsynchronized, metaphase control and KD cells were scored for spindle phenotype as described above for WT, KO-1 and KO-2 cells. Consistent with KO-1 and KO-2 cells, Myo10 KD cells exhibited a 20.7% reduction in the frequency of bipolar spindles and a 4.1-fold increase in the frequency of multipolar spindles (Fig. 1J). We conclude, therefore, that this core Myo10 KO phenotype is replicated in HeLa cells where Myo10 is transiently depleted using RNAi.

### MEFs isolated from a Myo10 conditional KO mouse and transduced with lenti-Cre phenocopy Myo10 KO HeLa cells

We recently described the creation of a Myo10 conditional knockout (cKO) mouse (tm1c) and several phenotypes exhibited by this mouse following its cross with a global Cre-deleter strain [13]. To provide additional support for the results obtained above using Myo10 KO HeLa cells, we treated primary MEFs isolated from this Myo10 cKO mouse with lentivirus expressing GFP-tagged Cre recombinase. Western blots showed that Myo10 was essentially undetectable 24 to 48 hours after lenti-cre transduction (Fig. 1C and Fig. S1C). Given this, non-transduced cKO MEFs and cKO MEFs 48 hours after transduction with lenti-Cre were scored for their metaphase spindle phenotype as described above for WT, KO-1 and KO-2 cells. As with Myo10 KO HeLa cells, scoring of >300 cells per line over three independent experiments showed that the lenti-Cre-transduced cKO MEFs exhibit a 35.6% decrease in the frequency of bipolar spindles, a 4.7-fold increase in the frequency of semipolar spindles, a 2.9-fold increase in the frequency of multipolar spindles, and a 3.6-fold increase in the frequency of non-bipolar spindles (semi-polar plus multipolar) (Fig. 3A). Also like Myo10 KO HeLa cells, optical sectioning of ∼200 metaphase cells exhibiting just two poles showed that the average distance separating the two spindle poles in Z was significantly greater in the lenti-Cre-transduced cKO MEFs than in the non-transduced cKO MEFs (Fig. 3B; 3C shows the distributions of these distances in 1 µm intervals). Finally, like Myo10 KO HeLa cells, the percentage of lenti-Cre-transduced cKO MEFs exhibiting chromosome misalignment and/or lagging chromosomes was 6.9-fold higher than for non-transduced cKO MEFs (Fig. 3D). We conclude, therefore, that cKO MEFs subjected to the conditional and rapid depletion of Myo10 largely phenocopy Myo10 KO HeLa cells as regards increased spindle multipolarity, abnormal spindle orientation relative to the substratum, and increased frequencies of chromosome misalignment and lagging chromosomes between metaphase and telophase.

**Figure 3.**
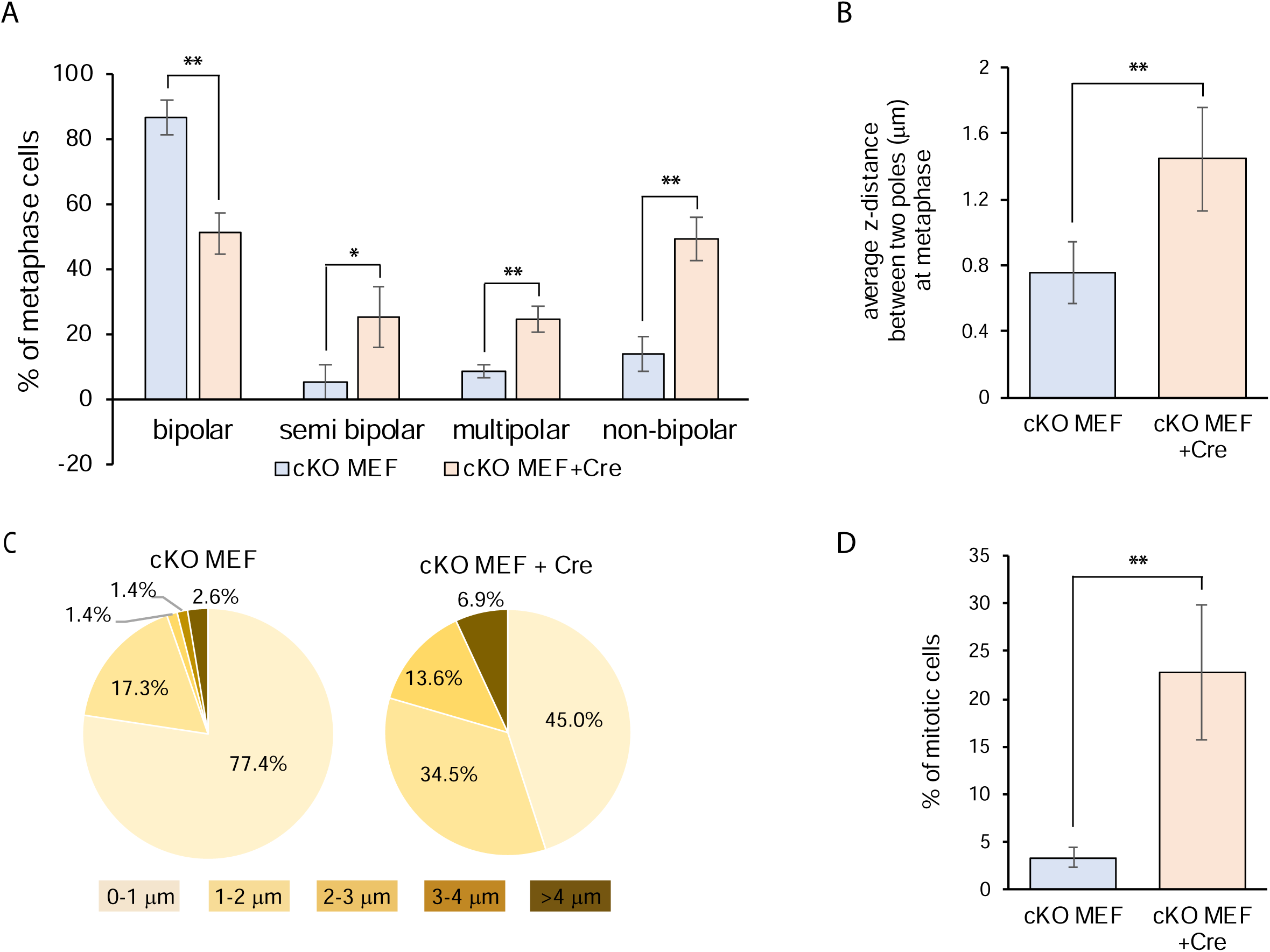
cKO MEFs transduced with lenti-Cre phenocopy Myo10 KO HeLa cells. (A) Quantitation at metaphase of the percent of bipolar, semi-bipolar, multipolar, and non-bipolar spindles in non-transduced cKO MEFs and cKO MEFs 48 hrs after lenti-cre transduction (159 cKO MEFS and 153 cKO MEFS plus Cre from 3 experiments). (B) Average distance in Z between the two poles of non-transduced cKO MEFs and cKO MEFs 48 hrs after lenti-cre transduction at metaphase (191 cKO MEFs and 208 cKO MEFs plus Cre over 3 experiments). (C) The distributions of the values in (B) in 1 um intervals. (D) Percent of mitotic, non-transduced cKO MEFs and cKO MEFs 48 hrs after lenti-cre transduction that possess misaligned and/or lagging chromosomes (169 cKO MEFs and 168 cKO MEFs plus Cre from 3 experiments).

### Myo10 KO MEFs from the straight Myo10 KO mouse exhibit an increase in the frequency of multipolar spindles whose magnitude depends on the phenotype of the embryo from which they are isolated and the density at which they are cultured

We next characterized the mitotic phenotype of MEFs isolated from the straight Myo10 KO mouse (tm1d; created by crossing the cKO Myo10 KO mouse (tm1c) with a global cre deleter strain;[13]). As described previously, the fate of embryos in single litters from the straight Myo10 KO mouse range from early embryonic lethality associated with exencephaly (a severe defect in neural tube closure) to viable mice exhibiting small body size, webbed digits, white belly spots, and microphthalmia. Given that the tm1d mouse is on a pure B6 background, this wide variation in embryo fate must be due to incomplete penetrance/variable expressivity rather than to variations between embryos in the inheritance of genetic modifiers [61] (although the non-genetic variations that dictate the fate of Myo10 KO embryos are unknown). We decided, therefore, to characterize MEFs isolated from both non-exencephalic and exencephalic tm1d embryos. We also decided to score the mitotic defects exhibited by these MEFs 24 hours after seeding them densely (5 × 10^4^ cells/cm^2^), at moderate density (2.5 × 10^4^ cells/cm^2^), and sparsely (0.5 × 10^4^ cells/cm^2^). This decision was based on measurements of growth rates for cells seeded at these three densities (referred to below and in the figures as “dense or D”, “moderate or M”, and “sparse or S”), where the growth of WT MEFs was unaffected by seeding density, while the growth of both non-exencephalic and exencephalic Myo10 KO MEFs was significantly slower at the sparse seeding density (Fig. S2A), as well as significantly slower than WT MEFs at the sparse seeding density (Fig. S2B). This result raised the possibility that Myo10 KO MEFs exhibit higher frequencies of mitotic defects and subsequent aneuploidy/cell senescence when provided with fewer spatial cues from neighboring cells.

To look for evidence that growth at lower cell densities does in fact increase the frequency of mitotic errors in Myo10 KO cells, we scored the frequency of multipolar spindles at metaphase in WT MEFs, non-exencephalic Myo10 KO MEFs, and exencephalic Myo10 KO MEFs 24 hours after seeding them at the three cell densities. While WT MEFs and non-exencephalic Myo10 KO MEFs were unaffected by cell density, exencephalic Myo10 KO MEFs exhibited higher frequencies of multipolar spindles as the cell density was lowered (Fig. S2C). Moreover, the frequencies for both KO MEFs were significantly higher than the frequency for WT MEFs at both moderate and sparse seeding densities (Fig. S2D). Additionally, the frequency exhibited by exencephalic Myo10 KO MEFs was significantly higher than the frequency exhibited by non-exencephalic Myo10 KO MEFs at both moderate and sparse seeding densities (Fig. S2D). Together, these results show that MEFs isolated from the straight Myo10 KO mouse exhibit a significant increase in the frequency of multipolar spindles, and that the severity of this defect depends on the phenotype of the embryo from which the MEFs are isolated, and on the density at which the MEFs are cultured. Finally, both non-exencephalic and exencephalic Myo10 KO MEFs grown at moderate density exhibit an increase in the frequency of spindles that are not parallel to the substrate (Fig. S2E and S2F). These results provide further support for a defect in cell adhesion during mitosis when Myo10 is missing.

### The multipolar phenotype exhibited by Myo10 KO HeLa cells is rescued by re-expression of Myo10

The fact that the multipolar spindle phenotype exhibited by Myo10 KO HeLa cells is also seen in Myo10 KD HeLa cells, Myo10 cKO MEFs after cre expression, and straight Myo10 KO MEFs argues that it is indeed a consequence of Myo10 depletion/loss. That said, we thought it was important to show that the re-introduction of Myo10 into Myo10 KO HeLa cells rescues the multipolar spindle phenotype before proceeding with efforts to define the mechanism(s) underlying this phenotype. Towards that end, we transfected KO-1 and KO-2 with mScarlet-Myo10 and scored mScarlet-Myo10-expressing cells at metaphase for spindle phenotype 24 hours post-transfection. Statistical analyses showed that Myo10 re-expression restored the values for the percentage of metaphase cells with bipolar and multipolar spindles in both KO lines to WT HeLa levels (Fig. 1K). Moreover, combining the data for semi-bipolar and multipolar spindles into non-polar spindles showed that both KO lines were completely rescued by Myo10 re-expression (Fig. 1L). Together, these results paved the way for subsequent efforts described below to define the mechanism(s) underlying the multipolar phenotype.

### Myo10 localizes dramatically in metaphase cells to the tips of retraction fibers and dorsal filopodia but rarely if at all to spindle poles and never within subcortical zones at the cell equator

We used HeLa KO-1 cells rescued with mScarlet-Myo10 to re-examine the localization of Myo10 in interphase and metaphase cells, focusing in particular on its reported localization in metaphase cells at spindle poles [42] and in subcortical actin clouds at the cell equator [43]. As expected [2-5], mScarlet-Myo10 localized robustly at the tips of both ventral filopodia (Fig. 4A1- A6) and dorsal filopodia (Fig. 4A7, A8; apical plane) during interphase, and at the tips of dorsal filopodia during metaphase (Fig. 4B1-B4; equatorial plane). Also as expected [43, 45], mScarlet-Myo10 localized dramatically at the tips of retraction fibers during metaphase (Fig. 4B5-B8). In contrast, Z-Stacks of 77 mScarlet-Myo10 expressing metaphase cells stained for DNA and F-actin showed no evidence that mScarlet-Myo10 localizes to spindle poles (see Z-Stack Movie 7 for a representative example). Consistently, Z-Stacks of 166 out of 171 mScarlet-Myo10 expressing metaphase cells stained for DNA and y-tubulin showed no evidence that the expressed myosin localizes to spindle poles (Fig. 4C1, C2; Z-Stack Movie 8). Finally, Z-Stacks of the 77 mScarlet-Myo10 expressing metaphase cells stained for DNA and F-actin, and the 171 mScarlet-Myo10 expressing metaphase cells stained for DNA and y-tubulin, showed no evidence the expressed myosin is enriched in equatorial, subcortical zones during metaphase (see Z-Stack Movies 7 and 8 for representative examples). So, with the exception of a tiny fraction of spindle poles, and excluding the diffuse fluorescence seen around spindles in some cells and in the cytoplasm of every cell (corresponding to freely-diffusing, cargo-free Myo10 [62-64]), mScarlet-Myo10 localized exclusively during metaphase to the tips of retraction fibers and dorsal filopodia.

**Figure 4.**
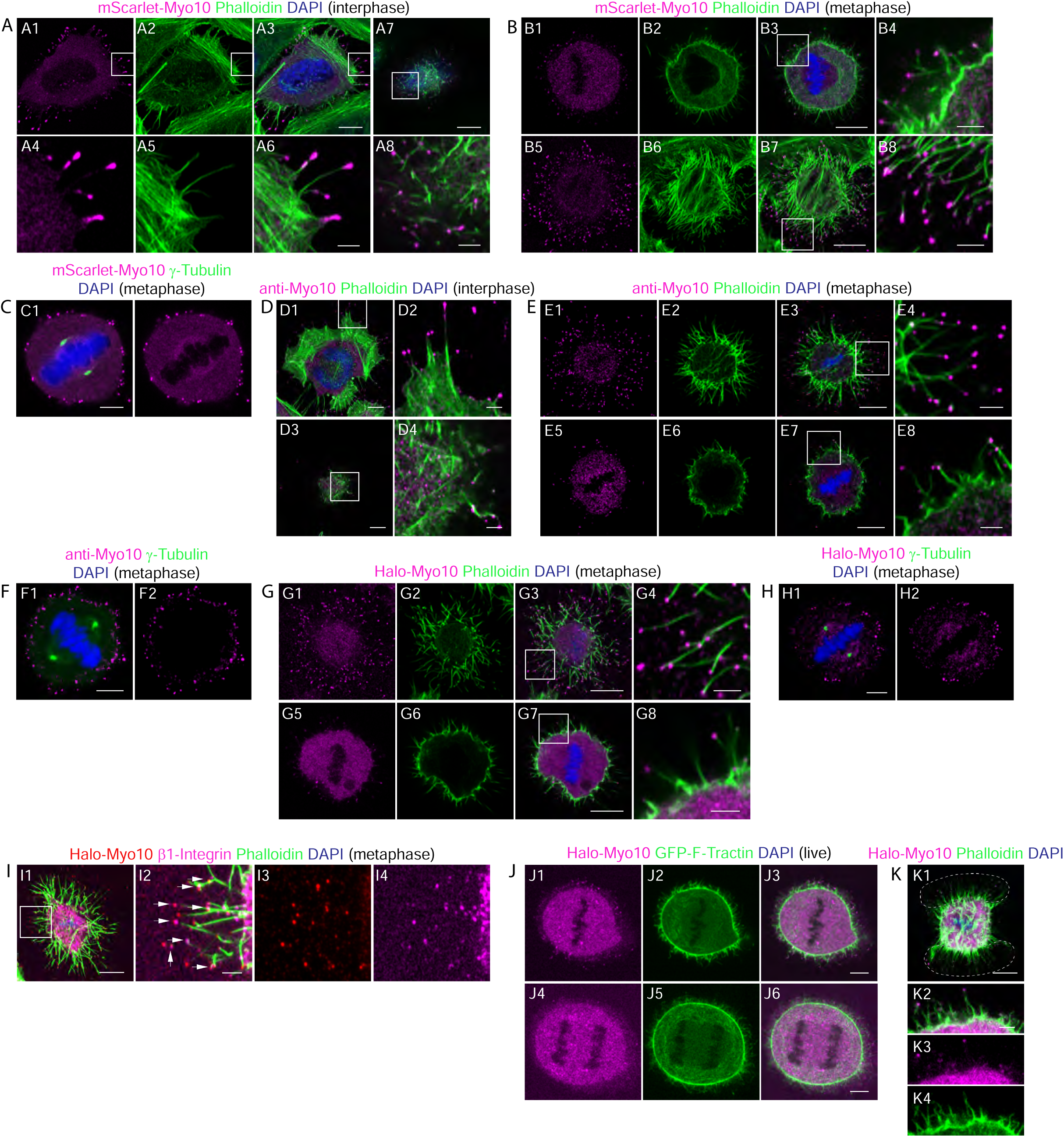
Myo10 localizes exclusively in metaphase cells to the tips of retraction fibers and dorsal filopodia. (A1-A6) An interphase Myo10 KO-1 cell expressing mScarlet-Myo10 that was fixed and stained with Alexa488-labeled Phalloidin and DAPI and imaged to show the accumulation of mScarlet-Myo10 at the tips of ventral filopodia. (A7, A8) Same as in A1-A6 except an apical section to show the accumulation of mScarlet-Myo10 at the tips of dorsal filopodia. (B1-B4) A metaphase Myo10 KO-1 cell expressing mScarlet-Myo10 that was fixed and stained as in A1-A8 and imaged to show the dorsal, mScarlet-Myo10-positive filopodia at the cell’s equator. (B5-B8) Same as B1-B4 except a ventral section to show the accumulation of mScarlet-Myo10 at the tips of retraction fibers (see also Z-Stack Movie 7). (C1, C2) A metaphase Myo10 KO-1 cell expressing mScarlet-Myo10 that was fixed and stained with anti-γ-tubulin-AF488 and DAPI (equatorial section; see also Z-Stack Movie 8). (D1-D4) An interphase HeLa cell that was fixed and stained with anti-Myo10, Alexa488-labeled Phalloidin and DAPI and imaged to show the accumulation of endogenous Myo10 at the tips of ventral (D1, D2) and dorsal (D3, D4) filopodia. (E1-E4) A metaphase HeLa cell that was fixed and stained as in D1-D4 and imaged to show the accumulation of endogenous Myo10 at the tips ventral retraction fibers. (E5-E8) Same as E1-E4 except an equatorial section to show the accumulation of endogenous Myo10 at the tips dorsal filopodia (see also Z-Stack Movie 9). (F1, F2) A metaphase HeLa cell that was fixed and stained with anti-Myo10, anti-γ-tubulin-AF488 and DAPI (equatorial section; see also Z-Stack Movie 10). (G1-G4) A metaphase Halo-Myo10 KI cell that was fixed and stained with Alexa488-labeled Phalloidin and DAPI and imaged to show the accumulation of Halo-Myo10 at the tips of ventral filopodia. (G5-G8) Same as G1-G4 except an equatorial section to show the accumulation of endogenously tagged Myo10 at the tips dorsal filopodia (see also Z-Stack Movie 11). (H1, H2) A metaphase Halo-Myo10 KI cell that was fixed and stained with anti-γ- tubulin-AF488 and DAPI (equatorial section; see also Z-Stack Movie 12). (I1-I4) A metaphase Halo-Myo10 KI cell that was fixed and stained Alexa488-labeled Phalloidin, DAPI and an antibody to the open, active form of β1 integrin (ITGB1 9EG7), and imaged to show the signals for Myo10 and active integrin in ventral retraction fibers (arrowheads points sites of colocalization between Myo10 and active integrin). (J1-J3) Single frame from a time lapse movie of a metaphase Halo-Myo10 KI cell expressing GFP-F-Tractin (equatorial section). (J4-J6) Same as J1-J3 except at anaphase (see also Movie 13). (K1) A Halo-Myo10 KI cell that was plated on an I bar pattern of fibronectin (dashed white outline) and fixed/stained at metaphase with Alexa488-labeled Phalloidin and DAPI (shown is a ventral section to reveal the distribution of retraction fibers; see also Z-Stack Movie 14). (K2-K4) Equatorial section of the cell in K1. Note that the bright dots in the Myo10 channel at the very perimeter of the metaphase cells stained for γ-tubulin in C, F and H (equatorial sections) are mScarlet-Myo10 (C), endogenous Myo10 (F) and Halo-Myo10 (H) at the tips of dorsal filopodia. The mag bars in A3, A7, B3, B7, D1, D3, E3, E7, G3, G7, I1 and K1 are 10 µm. The mag bars in C1, F1, H1, J3 and J6 are 5 µm. The mag bars in A6, A8, B4, B8, D2, D4, E4, E8, G4, G8, I2 and K2 are 2 µm.

To seek additional support for these findings, some of which are at odds with previous reports, we examined Myo10 localization in WT HeLa cells by staining for endogenous Myo10 using a Myo10 antibody whose specificity we confirmed using Myo10 KO cells (Fig. S3A1-A5, B1-B5). As expected, endogenous Myo10 localizes robustly in interphase cells at the tips of ventral and dorsal filopodia (Fig. 4D1-D4). Consistent with the results obtained using Myo10 KO cells rescued with mScarlet-Myo10, endogenous Myo10 localizes robustly in metaphase cells at the tips of retraction fibers (Fig. 4E1-E4) and dorsal filopodia (Fig. 4E5-FE; equatorial plane). Also consistent with the results obtained using mScarlet-Myo10 expressing KO cells, Z-Stacks of 83 metaphase cells stained for Myo10, DNA and F-actin showed no evidence that endogenous Myo10 localizes to spindle poles (see Z-Stack Movie 9 as a representative example). Consistently, Z-Stacks of 102 out of 102 metaphase cells stained for Myo10, DNA and y-tubulin showed no evidence that endogenous Myo10 localizes to spindle poles (Fig. 4F1, F2; Z-Stack Movie 10). Finally, Z-Stacks of the 83 metaphase cells stained for Myo10, DNA and F-actin, and the 102 metaphase cells stained for Myo10, DNA and y-tubulin, showed no evidence that endogenous Myo10 is enriched in equatorial, subcortical zones during metaphase (see Z-Stack Movies 9 and 10 for representative examples).

Given that the localization of Myo10 at spindle poles and in subcortical actin clouds at the equator of dividing cells were central to previous models for how Myo10 stabilizes spindle poles [42] and facilitates the clustering of supernumerary centrosomes [43], we sought to confirm our localization data by endogenous tagging of Myo10 using CRISPR (see Methods). This effort resulted in the creation of a Halo-Myo10 knockin (KI) HeLa cell line in which the Myo10 present in one allele is tagged at its N-terminus with Halo (Fig. S4A). As expected, Halo-Myo10 localizes robustly at the tips of filopodia in interphase cells (Fig. S4B1, B2) and at the tips of retraction fibers (Fig. 4G1-G4) and dorsal filopodia (Fig. 4G5-G8; equatorial plane) in metaphase cells. Consistent with the role that retraction fibers play in cell adhesion during mitosis, with Myo10’s ability to promote adhesion vis its FERM domain-dependent interaction with β1-integrin, and with the increase in the frequency of spindles that are not parallel to the substratum in Myo10 KO cells (Fig. 2), a significant fraction of Halo-Myo10 at the tips of retraction fibers colocalizes with the open, actin form of β1-integrin (Fig. 4I1-I4; see arrowheads). Importantly, Z-Stacks of 72 metaphase KI cells stained for DNA and F-actin showed no evidence that Halo-Myo10 is enriched at spindle poles (see Z-Stack 11 Movie for a representative example). Consistently, Z-Stacks of 107 out of 107 metaphase KI cells stained for DNA and y-tubulin showed no evidence that Halo-Myo10 localizes to spindle poles (Fig. 4H1, H2; Z-Stack Movie 12). Moreover, Z-Stacks of the 72 metaphase KI cells stained for DNA and F-actin, and the 107 metaphase KI cells stained for DNA and y-tubulin, showed no evidence that Halo-Myo10 is enriched in equatorial, subcortical zones during metaphase (see Z-Stack Movies 11 and 12 for representative examples). Consistently, time lapse movies of Halo-Myo10 KI cells expressing the F-actin probe GFP-F-Tractin failed to detect an enrichment of Halo-Myo10 within subcortical areas at the equator of either metaphase cells (Fig. 4J1-J3) or anaphase cells (Fig. 4J4-J6; see Movie 13). Regarding this latter localization, Kwon et al [43] argued that the Myo10 present there serves to connect the base of retraction fibers to the tips of astral microtubules to promote supernumerary centrosome clustering. To provide further support for this idea, they imaged cells dividing on an I bar pattern of fibronectin to restrict the attachment of retraction fibers to opposite sides of the cell cortex. In their images, over-expressed, GFP-tagged Myo10 was enriched within these two areas relative to the rest of the equatorial cortex.

We repeated this experiment using our Halo-Myo10 KI cells and saw no such enrichment (Fig. 4K1-K4; Z-Stack Movie 14; representative of ∼20 cells imaged). Similar results were seen with KO-1 cells expressing mScarlet-Myo10 (Z-Stack Movie 15). In summary, our images of Myo10 KO cells expressing moderate levels of mScarlet-Myo10, of WT cells immunostained for endogenous Myo10, and of endogenously tagged Myo10 in Halo-Myo10 KI cells all agree on three aspects of Myo10 localization during metaphase. The first is that Myo10 is seen rarely if at all at spindle poles. The second is that Myo10 is not concentrated in subcortical areas at the equator of dividing cells. The third is that Myo10 localizes dramatically at the tips of retraction fibers and dorsal filopodia. Together, these results suggest that Myo10 may support spindle pole integrity by acting at a distance, and that it may facilitate the clustering of supernumerary centrosomes by supporting cell adhesion during mitosis.

### Multipolar spindles in Myo10 KO HeLa cells arise from a combination of PCM fragmentation and an inability to cluster supernumerary centrosomes

We used our Myo10 KO HeLa cells to search for the cause(s) of spindle multipolarity when Myo10 is depleted. Four distinct cellular defects can give rise to multipolar spindles (reviewed in [65]). Two of these defects, cytokinesis failure and centriole overduplication, give rise to multipolar spindles because they generate interphase cells with extra centrosomes. The third defect, centriole disengagement, gives rise to multipolar spindles because both separated centrioles can serve as MTOCs. Finally, PCM fragmentation gives rise to multipolar spindles because the acentriolar PCM fragments that are generated can serve as MTOCs. Consistent with the results of Kwon et al [43] using Myo10 KD HeLa cells, we did not see a significant difference between WT and Myo10 KO HeLa cells in the average number of nuclei per cell (Fig. S5A1, S5A2), arguing that cytokinesis failure does not contribute significantly to their multipolar spindle phenotype. Centriole overduplication also does not appear to be driving the multipolar spindle phenotype, as unsynchronized WT and Myo10 KO HeLa cells have the same number of centrioles (Fig. S5B). For normal cells, which possess one centrosome, that would leave centriole disengagement and PCM fragmentation as the remaining possible causes of spindle multipolarity. For cancer cells, which often possess supernumerary centrosomes, the inability to cluster the extra spindle poles that ensue represents another possible cause of spindle multipolarity (reviewed in [57-59]). Given that a significant fraction of HeLa cells possess supernumerary centrosomes (see below), and that Myo10 has been implicated in promoting the clustering of the extra poles that ensue [43], subsequent experiments were designed to distinguish between centriole disengagement, PCM fragmentation, and an inability to cluster extra poles in cells with supernumerary centrosomes as causes of spindle multipolarity.

To accomplish this, we stained Myo10 KO HeLa cells for γ-tubulin and the centriole marker centrin-1 (along with DAPI), as this staining regimen reveals MTOCs arising from both acentriolar PCM fragments and centriole-containing structures. Imaging of unsynchronized, metaphase Myo10 KO HeLa cells stained in this way revealed examples of multipolar spindles caused by all three mechanisms discussed above. For example, Z-Stack Movie 16, along with the enlarged images of this cell’s three presumptive spindle poles (Fig. 5A1), represents an example of spindle multipolarity driven by centriole disengagement (note that two of this cell’s three presumptive spindle poles possess only one centriole). Z-Stack Movie 17, along with the enlarged images of this cell’s four presumptive spindle poles (Fig. 5A2), represents an example of spindle multipolarity driven by PCM fragmentation (note that two of this cell’s four presumptive spindle poles correspond to γ-tubulin-positive acentriolar foci). More extreme examples of multipolar spindles driven by PCM fragmentation were also seen (Fig. S6A and Z-Stack Movie 18; Fig. S6B and Z-Stack Movie 19). Finally, the still images in Figure 5, Panel A3, show an example of spindle multipolarity driven by a failure to cluster the extra spindle poles that arise in cells with supernumerary centrosomes (i.e. referred to below as centrosome de-clustering; note that this cell contains four presumptive spindle poles, each possessing two centrioles). Examples of partial supernumerary centrosome de-clustering were also observed (Fig. 5A4; note that only two of this cell’s three presumptive spindle poles (the two with four centrioles) were created by supernumerary centrosome clustering).

**Figure 5.**
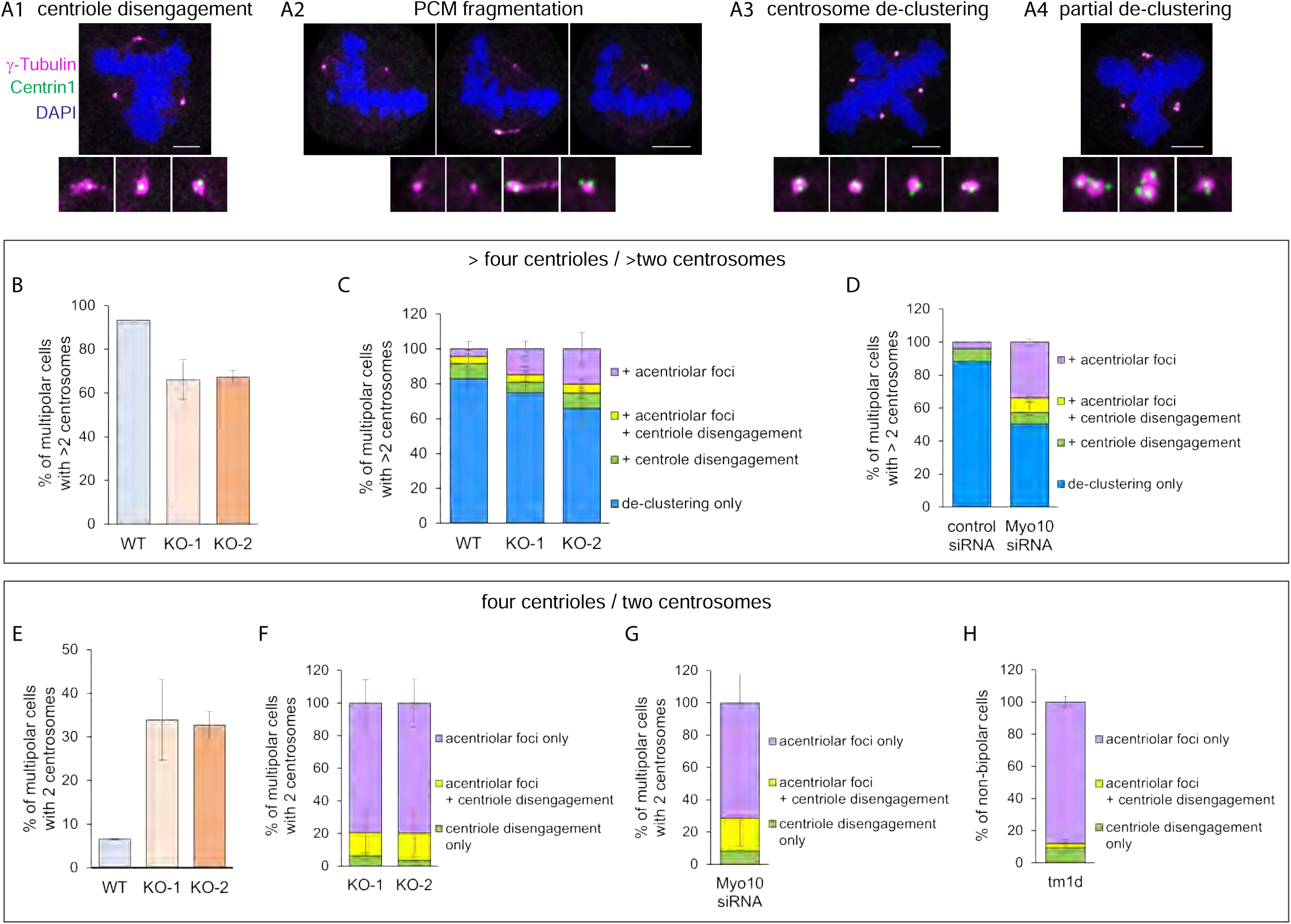
Causes of spindle multipolarity in Myo10-depleted HeLa cells and Myo10 KO MEFs. (A1-A4). Representative images of cells stained for γ-tubulin, centrin-1 and DNA that exhibit centriole disengagement (A1), PCM fragmentation (A2; shown are three different focal planes in one cell), centrosome de-clustering (A3), and partial centrosome de-clustering (A4). (B) Percent of multipolar WT HeLa, KO-1 and KO-2 that have more than 4 centrioles/2 centrosomes. (C) Percent of multipolar WT HeLa, KO-1 and KO-2 with more than 4 centrioles/2 centrosomes that exhibit centrosome de-clustering only (blue), centrosome de-clustering plus acentriolar foci (purple), centrosome de-clustering plus centriole disengagement (green), or centrosome de-clustering plus acentriolar foci and centriole disengagement (yellow). (D) Percent of multipolar HeLa cells exhibiting the mitotic defects described in Panel C that had been treated with control siRNA or Myo10 siRNA and that had more than 4 centrioles/2 centrosomes. (E) Percent of multipolar KO-1 and KO-2 with 4 centrioles/2 centrosomes. (F) Percent of multipolar WT HeLa, KO-1 and KO-2 with 4 centrioles/2 centrosomes that exhibit acentriolar foci (purple), acentriolar foci plus centriole disengagement (yellow), or centriole disengagement only (green). (G) Percent of multipolar HeLa cells exhibiting the mitotic defects described in Panel F that had been treated with Myo10 siRNA-treated and that had 4 centrioles/2 centrosomes (H) Percent of non-bipolar (semipolar plus multipolar) Myo10 KO MEFs isolated from the tm1d mouse that exhibit the mitotic defects described in Panel F. All mag bars are 5 µm.

To quantitate the relative contributions made by centriole disengagement, PCM fragmentation, and centrosome de-clustering to the multipolar spindle phenotype, we divided WT and Myo10 KO HeLa cells exhibiting multipolar spindles into two groups: those with more than four centrioles (i.e. cells that possessed supernumerary centrosomes) and those with four centrioles (i.e. cells that did not possess supernumerary centrosomes). For WT Hela, the vast majority of cells with multipolar spindles (93.4 ± 0.1%) contained more than four centrioles (Fig. 5B). In 82.9 ± 16.3% of these cells, every γ-tubulin spot contained at least two centrioles (Fig. 5C; WT blue), indicating that centrosome de-clustering was solely responsible for their multipolar spindle phenotype. For the remaining 17.1% of multipolar WT HeLa with more than four centrioles, 8.5 ± 8.0% exhibited centrosome de-clustering plus centriole disengagement (one or more γ-tubulin spots containing only one centriole) (Fig. 5C; WT green), 4.3 ± 4.0% exhibited centrosome de-clustering plus acentriolar foci arising (one or more γ-tubulin spots containing no centrioles) (Fig. 5C; WT purple), and 4.3 ± 4.0% exhibited centrosome de-clustering plus both centriole disengagement and acentriolar foci (Fig. 5C; WT yellow).

For KO-1 and KO-2 cells, about two thirds of cells with multipolar spindles contained more than four centrioles (66.1 ± 9.2% for KO-1 and 67.3 ± 3.0% for KO-2) (Fig. 5B). In roughly two thirds of these cells, every γ-tubulin spot contained at least two centrioles (74.9 ± 15.0% for KO- 1 and 65.9 ± 9.9% for KO-2) (Fig. 5C, KO-1 blue and KO-2 blue), indicating that centrosome de-clustering was mainly responsible for their multipolar spindle phenotype. Like WT HeLa, the remaining one third of multipolar KO cells with more than four centrioles exhibited centrosome de-clustering plus either centriole disengagement, acentriolar foci, or both (Fig. 5C; green, purple and yellow, respectively, for KO-1 and KO-2), although the percent of KO-1 cells and KO- 2 cells exhibiting acentriolar foci was 3.4-fold and 4.7-fold higher than in WT HeLa cells, respectively (Fig. 5C; compare the purple in KO-1 and KO-2 to the purple in WT). These results were supported by scoring those WT HeLa cells treated with Myo10 siRNA that contained more than four centrioles, which exhibited a 8.7-fold increase in the percentage of cells with acentriolar foci compared to HeLa cells treated with a control siRNA (Fig. 5D; compare the purple in Myo10 siRNA to the purple in control siRNA).

Finally, and most interestingly, were the results for cells exhibiting multipolar spindles that contained only four centrioles (i.e. cells that did not possess supernumerary centrosomes), which corresponded to about one third of Myo10 KO cells (33.9 ± 9.2% for KO-1 and 32.7 ± 3.0% for KO-2) but only a tiny fraction of WT HeLa cells (6.6 ± 0.1%) (Fig. 5E). Importantly, acentriolar foci arising from PCM fragmentation was by itself responsible for about ∼80% of the multipolar phenotype exhibited by this group of Myo10 KO cells (79.4 ± 14.3% for KO-1 and 79.6 ± 14.7% for KO-2) (Fig. 5F; purple in KO-1 and KO-2). Of the remaining ∼20% of multipolar KO cells with four centrioles, about three quarters exhibited acentriolar foci along with centriole disengagement (Fig. 5F; yellow in KO-1 and KO-2). In total, therefore, 93.6% of multipolar KO-1 cells lacking supernumerary centrosomes, and 96.4% of KO-2 cells lacking supernumerary centrosomes, exhibited acentriolar foci, arguing that PCM fragmentation is the primary cause of spindle multipolarity in these cells. Importantly, very similar results were obtained for HeLa cells subjected to transient Myo10 depletion using Myo10 siRNA. Specifically, 91.9% of multipolar Myo10 KD cells that did not possess supernumerary centrosomes exhibited acentriolar foci arising from PCM fragmentation (Fig. 5G; purple plus yellow in Myo10 siRNA). Together, these results argue that Myo10 supports spindle bipolarity in HeLa cells by promoting both PCM integrity and the clustering of supernumerary centrosomes.

### Multipolar spindles in Myo10 KO MEFs arise primarily from PCM fragmentation

Staining of MEFs isolated from the straight Myo10 KO mouse (tm1d), almost all of which possessed only four centrioles/two centrosomes at metaphase, showed that acentriolar foci alone were responsible for 88.0% of the multipolar spindles observed (Fig. 5H; purple). Of the remaining 12.0% of multipolar KO MEFs with four centrioles, about one third exhibited acentriolar foci plus centriole disengagement (Fig. 5H; yellow). In total, therefore, ∼91% of multipolar KO MEFs lacking supernumerary centrosomes exhibited acentriolar foci, arguing that PCM fragmentation is the primary cause of spindle multipolarity in these cells.

### Acentriolar foci arising from PCM fragmentation serve as MTOCs during mitosis

The preceding analyses of multipolar cells were done using unsynchronized cells stained for DNA (DAPI), centrioles (centrin-1) and γ-tubulin. Presumptive spindle poles were identified in these images based on DNA organization and on staining for γ-tubulin. In addition, over-exposure of the γ-tubulin signal often revealed evidence of microtubule asters emanating from these presumptive spindle poles. To provide additional evidence that PCM fragments lacking centrioles serve as spindle poles, we transfected Myo10 KO HeLa cells with dTomato-centrin-1 to mark centrioles, H2B-iRFP670 to mark chromatin, and EB1-EGFP to identify all MTOCs (i.e. those with centrioles and those lacking centrioles because they were created by PCM fragmentation). As expected, Myo10 KO HeLa cells containing two centriolar poles and undergoing normal bipolar mitoses exhibited EB1-EGFP comets emanating from two centriolar spindle poles (Fig. 6A1, A2 and Z-Stack Movie 20). Also as expected, Myo10 KO HeLa cells containing three centriolar poles and undergoing multipolar mitoses exhibited EBI-EGFP comets emanating from three centriolar spindle poles (Fig. 6B1, B2 and Z-Stack Movie 21). Importantly, imaging EBI-EGFP in Myo10 KO HeLa undergoing multipolar mitoses also revealed microtubule asters emanating from acentriolar poles in addition to centriolar poles (Fig. 6C1, C2 and Z-Stack Movie 22 enlarged images of centriolar and acentriolar spindle poles are shown in Movies 23 and 24, respectively). Together, these results confirm that PCM fragmentation creates acentriolar spindle poles that contribute to the multipolar phenotype exhibited by cells lacking Myo10.

**Figure 6.**
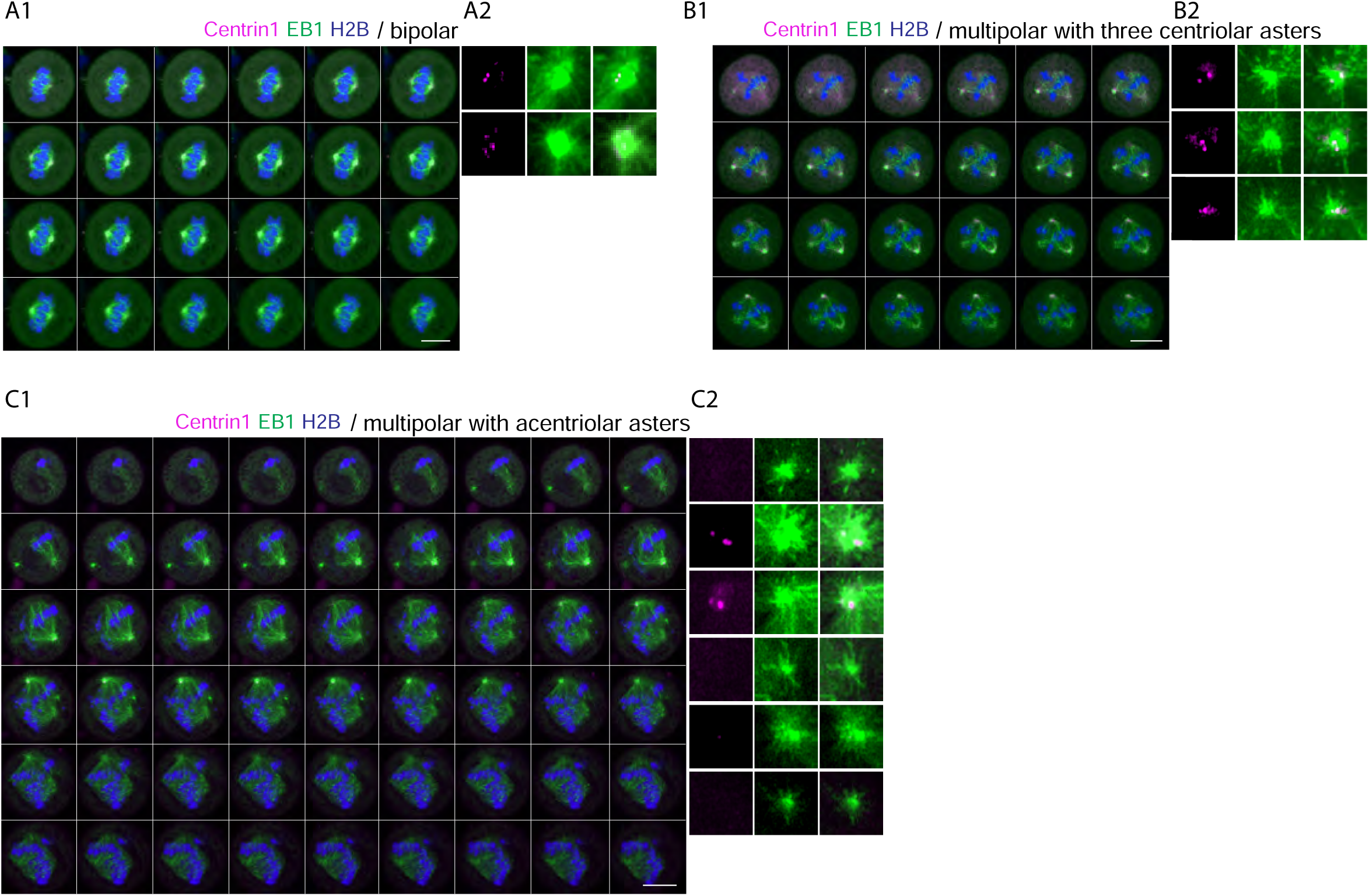
Acentriolar foci arising from PCM fragmentation serve as MTOCs during mitosis. (A1 and A2) Optical sections (A1) of a KO-1 cell containing two centriolar poles and undergoing a normal bipolar mitosis that had been transfected with EB1-EGFP to identify all MTOCs, dTomato centrin-1 to mark centrioles, H2B-iRFP670 to mark chromatin (see also Z-Stack Movie 20). The insets (A2) show enlargements of this cell’s two poles. (B1 and B2) Same as A1 and A2 except that this KO-1 cell contained three centriolar poles and was undergoing a multipolar mitosis (see also Z-Stack Movie 21). (C1 and C2) Same as A1 and A2 except that this KO-1 cell contained three acentriolar poles (i.e. negative for dTomato centrin-1) generated by PCM fragmentation in addition to three centriolar poles (see also Z-Stack Movie 22). Movies 23 and 24 show examples of EB1-EGFP comets emanating from a centriolar pole and an acentriolar pole, respectively. All the mag bars are 10 µm.

### PCM/pole fragmentation occurs primarily at metaphase, arguing that it is a force-dependent event

Defects in spindle pole integrity that lead to PCM/pole fragmentation are commonly revealed around metaphase when the chromosomal and spindle forces that the pole must resist increase [65]. To determine when the PCM fragments in Myo10 KO cells, we stained unsynchronized, mitotic KO cells for γ-tubulin, centrin-1 and DAPI. Scoring the percent of cells containing only two centrosomes that exhibited y-tubulin-positive, centriole-negative PCM fragments showed that these acentriolar fragments only begin to appear at prometaphase and peak in frequency at metaphase (Fig. 7A). Consistently, time lapse imaging of KO cells expressing dTomato-centrin-1, H2B-iRFP670, and EB1-EGFP reveals PCM/pole fragmentation and the formation acentriolar poles as they approach metaphase (see Movies 25). Time lapse imaging also shows that PCM/pole fragmentation at metaphase results in rearrangements of chromosomes that can lead to defects in chromosome segregation (see Movie 25). Together, these results indicate that cells do not enter mitosis with acentriolar PCM fragments, that PCM/pole fragmentation occurs primarily between prometaphase and metaphase, consistent with it being a force-dependent event, and that PCM/pole fragmentation results in defects in chromosome segregation.

**Figure 7.**
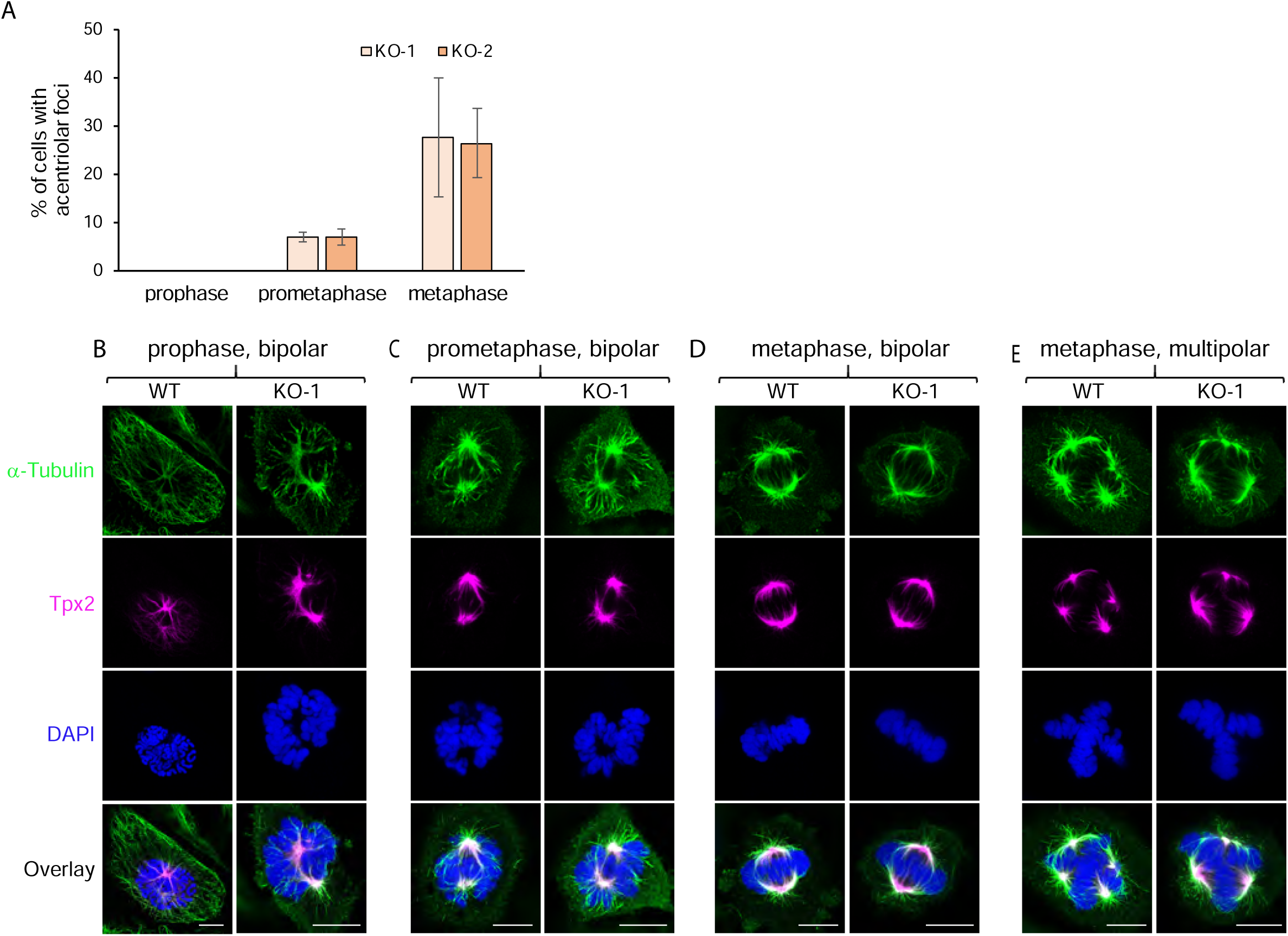
PCM/pole fragmentation is a force-dependent event that is not due to a defect in TPX2 localization. (A) Shown is the percent of unsynchronized KO-1 and KO-2 cells containing only two centrosomes that exhibited y-tubulin-positive, centriole-negative PCM fragments at prophase, prometaphase and metaphase. (B-E) Representative images of WT HeLa cells and Myo10 KO HeLa cells at prophase (B), prometaphase (C), and metaphase (D, E)) that were stained for α-tubulin, TPX2 and DNA (plus overlay) while undergoing bipolar divisions (B-D) or a multipolar division (E). All the mag bars are 10 µm.

### PCM fragmentation in Myo10 KO HeLa cells is not due to a defect in the pole localization of TPX2

As reviewed in the Introduction, Bement and colleagues [42] showed previously that embryonic frog epithelial cells depleted of Myo10 using morpholinos exhibit spindle pole fragmentation at metaphase, leading to an increase in the frequency of multipolar spindles. Moreover, they attributed this apparent defect in pole integrity to a defect in the localization of the spindle pole assembly factor TPX2 at and near poles when Myo10 is depleted. To determine whether the PCM fragmentation observed here is also caused by a defect in TPX2 localization, we stained WT and Myo10 KO HeLa cells for α-tubulin, TPX2 and DNA at prophase, prometaphase and metaphase. Figure 7, Panels B-D, show that KO cells undergoing bipolar mitosis do not exhibit any obvious defect in the localization of TPX2 at and near spindle poles at all three mitotic stages. Similarly, the localizations of TPX2 in WT and KO HeLa cells undergoing multipolar mitosis are indistinguishable (Fig. 7E). These results indicate that the PCM fragmentation observed here cannot be attributed to a defect in the recruitment of TPX2 to poles when Myo10 is missing.

### The defect in PCM/pole integrity in Myo10 KO HeLa cells is not due to an obvious defect in spindle pole maturation

Defects in spindle pole maturation can lead to defects in pole integrity [65]. To look for a possible defect in spindle pole maturation in Myo10 depleted cells, we stained WT, KO-1 and KO-2 HeLa cells at prophase, prometaphase and metaphase for centrin-1, DNA, and the pole protein CDK5Rap2, which is known to accrue in maturing poles where it recruits γ-TuRC to promote microtubule nucleation [66, 67]. Stained cells possessing only two poles were optically sectioned in 0.25 µm intervals and the total intensity of the CDK5Rap2 signal obtained by summing slices (see Methods for additional details). Representative images of stained poles in KO-1 cells at prophase, prometaphase and metaphase are shown on Figure 8, Panels A-C, respectively. Importantly, quantitation revealed no significant differences between WT cells and either KO-1 or KO-2 in the pole content of CDK5Rap2 at all three mitotic phases (Fig. 8D). This result, together with the fact that Myo10 is not present at spindle poles in Halo-Myo10 KI cells (Fig. 4), indicates that the defect in PCM integrity is not part of a general defect in pole maturation, and suggests that Myo10 promotes PCM integrity at a distance (see Discussion).

**Figure 8.**
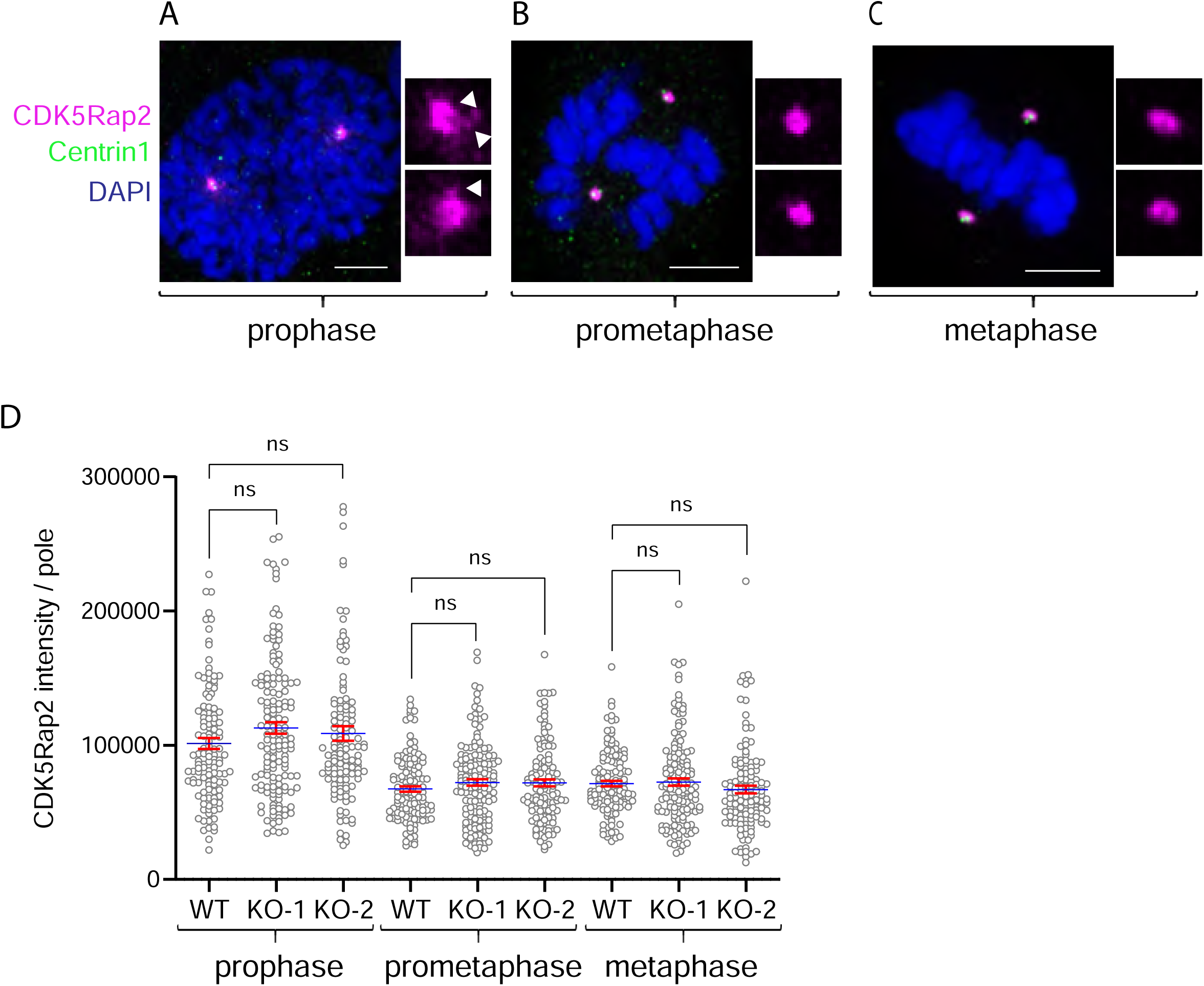
The defect in PCM/pole integrity in Myo10 knockout cells is not due to an obvious defect in spindle pole maturation. (A-C) Representative images of unsynchronized KO-1 cells stained at prophase (A), prometaphase (B), and metaphase (C) for centrin-1, DNA, and the pole protein CDK5Rap2. The enlarged insets show the signal for CDK5Rap2 only (the white arrowheads in the insets for (A) point to CDK5Rap2-postive pericentriolar satellites). (D) Quantitation of the amount of CDK5Rap2 at poles in WT HeLa, KO-1 and KO-2 at prophase, prometaphase and metaphase (125 to 150 cells per condition from 3 independent experiments). The fact that the pole content of CDK5Rap2 is higher at prophase than at prometaphase and metaphase may be due to the presence of CDK5Rap2-positive pericentriolar satellites surrounding the poles at prophase but not at prometaphase and metaphase. All mag bars are 5 µm.

### Complementation experiments show that Myo10 must interact with both integrins and microtubules to promote spindle pole integrity, and that its interaction with integrins is key to its ability to promote supernumerary centrosome clustering

We used complementation of HeLa Myo10 KO cells to access the contributions made by the myosin’s integrin-binding FERM domain and microtubule-binding MyTH4 domain to its ability to promote PCM/pole stability and the clustering of the extra poles that arise in cells with supernumerary centrosomes. To accomplish this, we introduced function blocking point mutations within these two domains in the context of full length, mScarlet tagged Myo10 (see Methods and Fig. S8 for details). WT Myo10 and these two mutated versions of Myo10 (referred to as mut-FERM Myo10 and mut-MyTH4 Myo10) were expressed in KO-1 cells by creating stable, Tet-inducible lines. As anticipated, doxycycline addition resulted in the expression of all three proteins with the correct molecular weight for tagged, full length Myo10 (Fig. 9A) and at a level that slightly exceeded that of endogenous Myo10 (WT Myo10, mut-FERM Myo10 and mut-MyTH4 Myo10 were expressed at about 2.2, 1.6 and 2.1 times the level of endogenous Myo10, respectively). Moreover, like WT Myo10 (Fig. 4; Fig. S7A1-A3 and D1- D4), both Myo10 mutants localize at the tips of interphase filopodia (Fig. S7B1-B3 and C1-C3) and metaphase retraction fibers (Fig. S7E1-E4 and F1-F4). Given these observations, an inability to rescue, whether partial or complete, cannot be attributed to lack of expression, significant variation between rescue constructs in expression level, or mislocalization of the mutant proteins.

**Figure 9.**
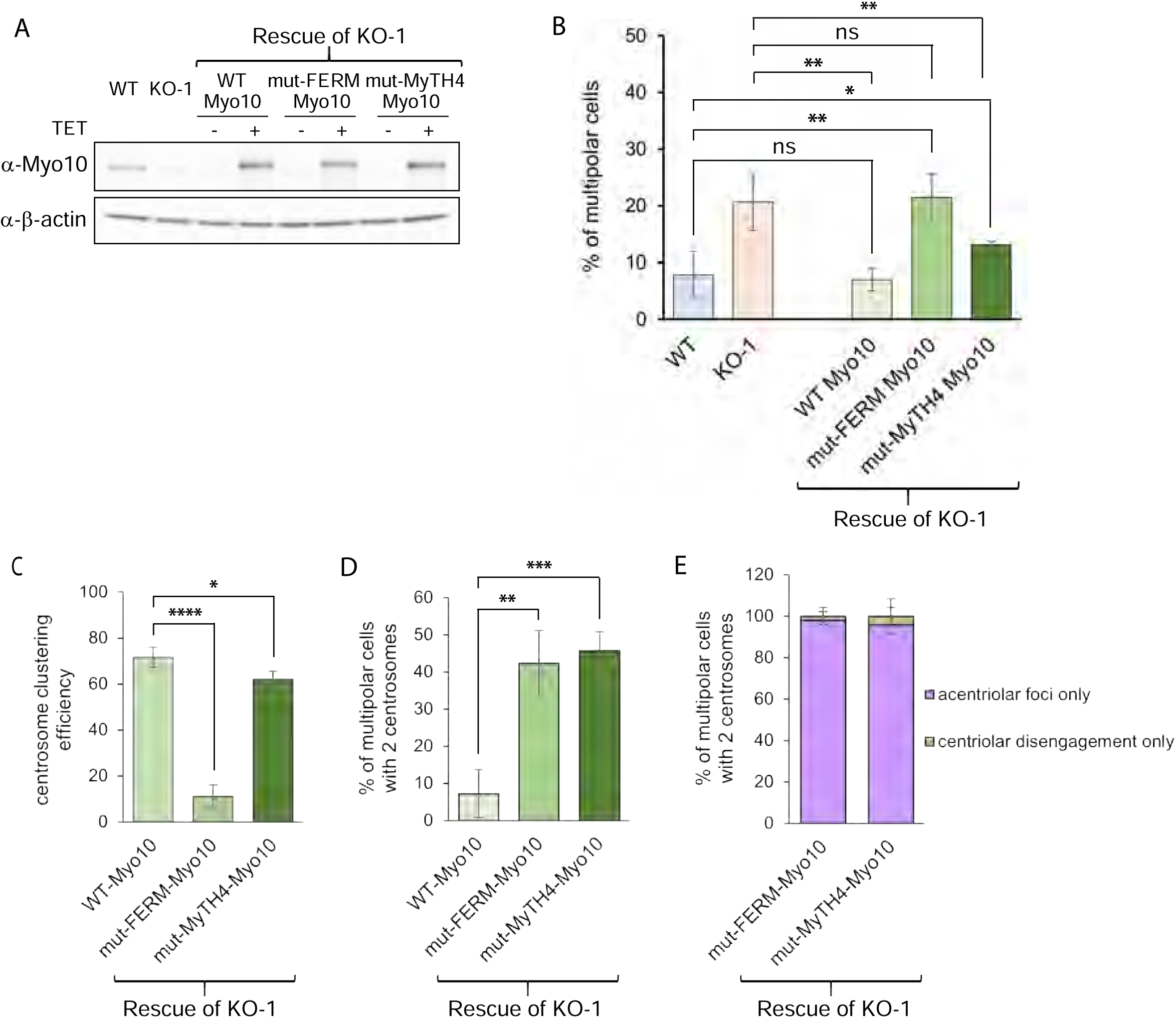
Complementation experiments reveal the contributions made by Myo10’s FERM and MyTH4 domains to spindle pole integrity and supernumerary centrosome clustering. (A) Westerns blots of whole cell extracts prepared from WT HeLa, KO-1, and KO-1 rescued with mScarlet tagged WT Myo10, mut-FERM Myo10 or mut-MyTH4 Myo10 plus/minus doxycycline addition, and probed for Myo10 and β-actin as a loading control (see Methods for details). (B) Quantitation at metaphase of the percent of multipolar spindles in WT HeLa, KO-1, and KO-1 rescued with WT Myo10 (338 cells), mut-FERM Myo10 (386 cells) or mut-MyTH4 Myo10 (335 cells) (from 3 experiments). (C) The efficiency of supernumerary centrosome clustering in KO-1 rescued with WT Myo10 (76 cells), mut-FERM Myo10 (69 cells) or mut-MyTH4 Myo10 (75 cells) (from 3 experiments). (D) The percent of multipolar cells with 2 centrosomes in KO-1 rescued with WT Myo10 (24 cells), mut-FERM Myo10 (44 cells) or mut-MyTH4 Myo10 (45 cells) (from 3 experiments). (E) The percent of multipolar KO-1 cells with 2 centrosomes rescued with mut-FERM Myo10 or mut-MyTH4 Myo10 that exhibited acentriolar foci only or centriolar disengagement only (from 3 experiments).

As expected, WT Myo10 fully rescued the multipolar phenotype of KO-1 cells (Fig. 9B). To estimate the contribution that the FERM and MyTH4 domains make to the clustering of extra spindle poles, we divided the number of rescued cells possessing more than four centrioles and a bipolar spindle by the total number of rescued cells with more than four centrioles (which encompass bipolar, semipolar and multipolar cells) to obtain a value for the efficiency of supernumerary centrosome clustering. While cells rescued with WT Myo10 yielded a value of 72 ± 4.4%, cells rescued with mut-FERM Myo10 yielded a value of 11 ± 4.8% (Fig. 9C). This result indicates that Myo10’s ability to interact with integrins is essential for its ability to promote supernumerary centrosome clustering. In contrast, cells rescued with mut-MyTH4 Myo10 yielded a value of 62 ± 3.4%, indicating that Myo10’s ability to interact with microtubules is not essential for its ability to promote supernumerary centrosome clustering (Fig. 9C). That said, the MyTH4 domain makes some contribution as cells rescued with mut-MyTH4 Myo10 were slightly less efficient at clustering extra poles than cells rescued with WT Myo10.

To estimate the contribution that each domain makes to the maintenance of PCM/pole stability, we determined the underlying cause of multipolarity in rescued cells possessing only four centrioles. This was a minute fraction of cells rescued with WT Myo10 but about half of the cells rescued with mut-FERM Myo10 or mut-MyTH4 Myo10 (Fig. 9D). For these latter two groups, multipolarity was associated almost entirely with PCM fragmentation, as indicated by the presence of acentriolar foci only (Fig. 9E). This result argues that Myo10’s FERM domain-dependent interaction with integrin and its MyTH4 domain-dependent interaction with microtubules are both required for its ability to promote PCM/pole integrity.

## Discussion

Here we showed that the primary driver of spindle multipolarity exhibited by Myo10 KO MEFs and by Myo10 KO HeLa cells lacking supernumerary centrosomes is PCM fragmentation, which creates y-tubulin-positive, centriole-negative microtubule asters that serve as additional spindle poles. We also showed that the primary driver of spindle multipolarity in HeLa cells possessing supernumerary centrosomes is an inability to cluster the extra spindle poles that ensue. Importantly, all of our scoring was done using unsynchronized cells, as treatments used to synchronize cells by delaying mitotic entry (e.g. low dose nocodazole) can lead to the formation of multipolar spindles with abnormal centriole distributions at poles (e.g. single or no centrioles) due to cohesion fatigue [65].

Extra spindle poles are created in the absence of centrosome amplification either through centriole disengagement or PCM fragmentation (reviewed in [65]). Here we showed for the first time that the appearance of extra spindle poles in cells lacking Myo10 (and not possessing supernumerary centrosomes) is due almost entirely to PCM fragmentation, i.e. that centriole disengagement makes only a minor contribution. This puts the focus on Myo10’s contribution to maintaining spindle pole integrity squarely on its contribution to maintaining PCM integrity. PCM fragmentation typically occurs when the physical integrity of spindle poles is compromised in some way, causing them to fragment when subjected to the strong chromosomal and spindle forces that ramp up during metaphase (reviewed in [65]). Consistently, the PCM fragmentation events we observed in Myo10 KO HeLa cells occurred primarily around metaphase. While we do not at present know the molecular mechanism by which Myo10 supports PCM integrity, we made several observations that can direct future work. First, we did not see any obvious role for Myo10 in pole maturation, as the pole localization of both TPX2 and a standard marker for pole maturation (CDK5Rap2) are normal in Myo10 KO HeLa cells. Second, we did not see Myo10 at spindle poles in either immunostained cells or Halo-Myo10 KI cells. So, with the exception of a tiny fraction of KO HeLa cells expressing mScarlet-tagged Myo10, we found no evidence that Myo10 localizes to spindle poles. Together, these results indicate that the loss of PCM integrity in Myo10 KO cells is not part of a general defect in pole maturation, and they suggest that Myo10 functions at a site other than the spindle pole to maintain PCM integrity. However Myo10 accomplishes this task, our complementation data showed that its microtubule-binding MyTH4 domain and its integrin-binding FERM domain are both required.

One possible explanation for how Myo10 might promote PCM integrity at a distance involves its ability regulate the CDK1 kinase inhibitor WEE1 [49]. Normally, CDK1 activity increases during prophase, is highest at metaphase, and then decreases at anaphase onset to promote mitotic progression [68]. Importantly, Bement and colleagues presented evidence that Myo10 binds WEE1 as cells approach metaphase to prevent WEE1 from inactivating CDK1, and then releases WEE1 at the end of metaphase so that it can suppress CDK1 activity and promote mitotic progression [49]. In this way, then, the interaction of Myo10 with WEE1 could coordinate Myo10’s role in attaining correct spindle orientation [42] with anaphase onset.

One interesting ramification of this model has to do with the role that CDK1 activity plays in determining the localization of the microtubule-binding and dynein-anchoring protein NuMA [68-71]. Normally, the robust activity of CDK1 up to metaphase keeps NuMA largely phosphorylated, which in turn keeps it largely localized to spindle poles, where it promotes pole stability and pole focusing, and in the spindle, where it promotes load sharing by crosslinking k-fibers to interpolar microtubules (although some unphosphorylated NuMA present in the cortex serves to promote metaphase spindle positioning by recruiting dynein) [68-72]. At anaphase onset, however, CDK1 activity drops. This drop, together with the action of the phosphatase PPP2CA, causes NuMA’s phosphorylation state to drop. This precipitates the translocation of significant amounts of NuMA from spindle poles and the spindle to the cortex, where it, together with cortex-bound dynein, drives spindle elongation during anaphase [68-72]. Given all this, one would predict that in the absence of Myo10-dependent WEE1 sequestration (i.e. in Myo10 KO cells), CDK1 would be inhibited prematurely, leading to the premature translocation of NuMA from spindle poles and the spindle to the cortex. Importantly, this might result in a reduction in PCM/pole stability [68, 69, 73] that, together with the premature escalation of dynein-dependent cortical pulling forces, could lead to PCM fragmentation at metaphase. Future efforts should examine NuMA’s dynamic localization across mitosis in cells lacking Myo10, and should seek to define the physiological significance of Myo10-WEE1 interaction by complementing Myo10 KO cells with a version of Myo10 that cannot bind WEE1. Finally, it remains possible that the signal for Myo10 in the spindle, while faint, diffuse and not always seen (Fig. 4), represents a pool of microtubule-associated Myo10 that promotes PCM integrity in some way. Notably, this mechanism might explain why Myo10 requires a functional MyTH4 domain to support PCM integrity.

While our data indicated that PCM fragmentation contributes significantly to the formation of multipolar spindles in Myo10 KO HeLa cells possessing supernumerary centrosomes, it indicated that the primary cause of spindle multipolarity in these cells is an inability to cluster the extra spindle poles that arise. This result confirms and extends the seminal observations made by Kwon and colleagues using RNAi screens for genes whose expression promotes supernumerary centrosome clustering, which implicated Myo10 in MDA-MB-231 cancer cells and fly Myo10A (a homolog of the human MyTH4/FERM myosin, Myo15) in near-tetraploid *Drosophila* S2 cells [48]. Importantly, this screen also identified the pole focusing, microtubule minus end-directed kinesin 14 family member HSET (Ncd in Drosophila) as being essential for supernumerary centrosome clustering. In subsequent work, Kwon and colleagues showed that Myo10 with a mutated MyTH4 domain cannot rescue the defect in supernumerary centrosome clustering exhibited by Myo10 knockdown cells overexpressing PLK4 [43]. Given this, and given their evidence that Myo10 present within subcortical actin clouds at the equator of dividing cells cooperates in a MyTH4 domain-dependent manner with dynein to position the metaphase spindle, Kwon et al proposed that this same pool of Myo10 cooperates in a MyTH4 domain-dependent manner with HSET to drive the clustering of supernumerary centrosomes.

We presented data here, on the other hand, that supports a FERM domain/adhesion-centric mechanism for how Myo10 promotes supernumerary centrosome clustering. First, we confirmed using a variety of approaches that Myo10 localizes dramatically at the tips of retraction fibers in mitotic HeLa cells. These actin-based structures have been considered for decades as being essential for the integrin-dependent adhesion of cells undergoing mitosis [50-55]. Consistently, the signals for endogenous Myo10 and open, active integrin were seen to overlap significantly within retraction fibers. Second, we showed that the spindle in Myo10 KO cells is often not parallel to the substratum. This result, and a similar result reported by Toyoshima and colleagues in Myo10 knockdown HeLa cells [44], is consistent with Myo10 depletion causing a defect in cell adhesion during mitosis. Third, and most importantly, our complementation experiments showed that Myo10’s ability to interact with integrins via its FERM domain is essential for its ability to promote supernumerary centrosome clustering. We note that all of these results are completely in line with the now strong evidence that Myo10 supports the formation of adhesions within filopodia and lamellipodia during interphase by virtue of its FERM domain-dependent interaction with β1-integrin [12, 23-25]. Moreover, they are in line with micropatterning data showing that the ability of cancer cells to cluster their supernumerary centrosomes is influenced significantly by the pattern of adhesion [48]. What our results are not in line with, however, is the MyTH4 domain-centric model proposed by Kwon et al for how Myo10 promotes supernumerary centrosome clustering [43]. First, none of our localization approaches, including the imaging of Halo-Myo10 KI cells undergoing division on an I bar pattern of ECM to focus retraction fiber: cortex interactions within a narrow region of the cortex, showed that Myo10 is concentrated in subcortical actin clouds at the equator of dividing HeLa cells. We suggest that Myo10 overexpression, combined with the collapse of Myo10-positive dorsal filopodia onto the cell surface upon fixation, may be responsible for the localization data presented by Kwon et al [43]. Second, our complementation data showed that Myo10’s ability to interact with microtubules via its MyTH4 domain made only a minor contribution to the myosin’s ability to promote supernumerary centrosome clustering. While it remains possible that the MyTH4 domain-centric mechanism proposed by Kwon et al [43] makes some contribution to supernumerary centrosome clustering, our results argue that Myo10’s ability to promote retraction fiber-based cell adhesion during mitosis via it FERM domain-dependent interaction with integrins plays the major role. As to how this would promote the clustering of extra spindle poles to create a bipolar spindle, we suggest that it serves as one of several anchors that together support the HSET-dependent microtubule forces driving pole focusing by maintaining spindle tension [56]. By extension, we hypothesize that Myo10’s FERM domain-dependent interaction with integrins is promoting the dynein-dependent positioning of the mitotic spindle in exactly the same way [45-47]. Finally, Myo10’s role in supporting cell adhesion during mitosis may have added importance given that conventional focal adhesions largely disappear as cell enter mitosis [55].

If not clustered, supernumerary centrosomes cause multipolar cell divisions that result in chromosome miss-segregation, aneuploidy and cell death [57-59]. Given this, and given that supernumerary centrosomes are rare in normal cells and common in cancer cells, drugs that inhibit the clustering of supernumerary centrosomes should selectively kill cancer cells. Indeed, compounds that inhibit HSET have been tested as anti-cancer therapeutics [74]. By the same token, a specific inhibitor of Myo10 might also serve to selectively kill cancer cells. Moreover, it might work synergistically with inhibitors of HSET. This seems likely given the results of Kwon et al [48], who showed using assays that quantified the suppression of supernumerary centrosome clustering that actin disassembly was not synergistic with Myo10 knockdown (presumably because they act in the same pathway), but was synergistic with HSET inhibition. Notably, several recent studies have provided additional support for the idea that inhibitors of Myo10 might be effective as cancer therapeutics. First, Collucio and colleagues [75] reported that depletion of Myo10 in mouse models of melanoma reduces melanoma development and metastasis and extends medial survival time. Similarly, Rosenfeld and colleagues [76] reported that the lifespan of mice with glioblastoma is extended significantly on a Myo10 KO background. Third, Zhang and colleagues [77] reported that the progression of breast cancer tumors in mice is inhibited by Myo10 depletion and accelerated by Myo10 overexpression. More generally, many cancer cell types exhibit elevated levels of Myo10, which may help support their growth by promoting the clustering of their extra centrosomes [40, 78, 79]. These and other studies [40], together with the data presented here, provide strong justification for performing screens to identify inhibitors of Myo10 as possible cancer therapeutics.

## Materials and Methods

### Cell culture and transfection

HeLa cells (ATCC, CCL-2), HEK-293T cells (ATCC, CRL-3216), and primary MEFs were cultured in high glucose DMEM (Gibco) supplemented with 10% FBS (Gibco) and Antibiotic-Antimycotic solution (Gibco), and were maintained at 37°C in a 5% CO_2_ humidified incubator. MEFs were prepared from ∼E19 embryos as described previously [80]. Only low passage number MEFs (<P3) were used for quantitating knockout phenotypes. For imaging purposes, cells were cultured in coverglass bottom chamber slides (Cellvis) coated with fibronectin (10 ng/ml; Gibco). Cells were transfected using either Lipofectamine 3000 (Invitrogen) or an Amaxa nucleofection apparatus (Lonza) according to the manufacturer’s instructions. To knock down Myo10, the ON-TARGETplus human Myo10 siRNA-SMARTpool from Dharmacon RNA Technologies (L-007217-00) was transfected into WT Hela cells using Lipofectamine RNAiMAX reagent (Invitrogen). To increase knockdown efficiency, the transfection was carried out twice with two-days interval in reverse fashion. The I bar-shaped fibronectin micropatterns were purchased from CYTOO and used according to the manufacturer’s instructions. All cells were checked routinely by PCR for possible contamination by mycoplasma.

### Mice

WT C57BL/6 mice were purchased from Jackson Laboratories (#000664). The creation of the straight Myo10 KO mouse (tm1d) and the Myo10 cKO mouse (tm1c) are described in our previous study [13]. Both Myo10 KO mice are on a pure B6 background. All animal experiments were approved by the Animal Care and Use Committee of the National Heart, Lung, and Blood Institute, in accordance with the National Institutes of Health guidelines.

### Antibodies and immunofluorescence

The following primary antibodies were used: mouse anti-γ-tubulin (MilliporeSigma, T6793, 1:300), mouse anti-α-tubulin (Abcam, ab7291, 1:300), rabbit anti-α-tubulin (Abcam,ab52866, 1:300), rabbit anti-myo10 (MilliporeSigma, HPA024223, 1:300), mouse anti-centrin1 (MilliporeSigma, 04-1624, 1:200), rabbit anti-centrin-1 (Abcam, ab101332, 1:200), rabbit anti-CDK5Rap2 (MilliporeSigma, 06-1398, 1:200), rabbit anti-pericentrin (Abcam, ab4448, 1:200), mouse anti-β-actin (Abcam, ab6276, 1:10,000), and mouse anti-Tpx2 (Santa Cruz Biotechnology, sc-53775, 1:200), rat-anti-CD29 (BD Pharmingen, 553715, 1:200). AlexaFluor-conjugated and HRP-conjugated secondary antibodies were purchased from Jackson ImmunoResearch Laboratories. DAPI, Alexa Fluor 488 labeled Phalloidin, and Alexa Fluor 568 labeled Phalloidin were purchased from ThermoFisher Scientific. Cells were fixed in 4% PFA for 10 min at RT unless subsequent staining was for centrosomes and pole-related proteins, in which case the cells were subjected to a two-step fixation method involving 1.5% PFA for 5 min followed by ice-cold MeOH for 5 min at −20°C. All PFA solutions were prepared in Cytoskeleton Stabilization Buffer (150 mM NaCl, 5 mM EGTA, 5 mM glucose, 5 mM MgCl_2_, and 10 mM PIPES (pH 6.8)). Fixed cells were permeabilized and blocked by incubation for 15 min in PBS containing 0.15% Saponin and 5% fetal bovine serum (FBS). Fixed, permeabilized cells were incubated at RT for 1 hr with primary and secondary antibodies diluted in blocking solution, with an intervening wash cycle using PBS.

### DNA constructs, mutagenesis of Myo10, and complementation

In Fusion HD cloning (Takara 638910) was used to switch full length WT mouse Myo10 in EGFP-C1 to mScarlet-C1, and to clone WT and mutant versions of mScarlet-Myo10 into an Xlone plasmid (Addgene plasmid # 96930). The Xlone-GFP backbone plasmid was linearized by treating with restriction enzymes SpeI-HF (NEB R3133L) and KpnI-HF (NEB R3142L). To create the rescued cell lines, HeLa Myo10 KO1 cells were nucleofected using an Amaxa nucleofection apparatus (Lonza; program I-013) with Piggybac plasmid and either WT Xlone-mScarlet-Myo10 or mutant Xlone-mScarlet-Myo10. Cells were subsequently plated in 3 wells of a 6-well plate and left to proliferate. At 90% confluency, cells were treated with 8 ug/ml of blasticidin (Gold Biotechnology, B-800-100) for 48 h to select for cells that had integrated the Xlone-mScarlet-Myo10 plasmid. Afterwards cells were left to proliferate in media without blasticidin, and 24 h prior to cell sorting were treated with 2 ug/ml of doxycycline (Sigma-Aldrich, D9891-5G). Cells positive for mScarlet fluorescence were sorted, propagated and used in the rescue experiments. mScarlet tagged mut-FERM Myo10 was created by changing Ile 2041, which resides within the F3 lobe of Myo10’s FERM domain, to Gln using PCR-based mutagenesis. The rationale for choosing this mutation to attenuate Myo10: integrin interaction is as follows: (i) the structure of the F3 lobe within the FERM domain of mouse talin 2 complexed with the cytoplasmic tail of human β1-integrin (PDB entry 3G9W), and the predicted structure of the complex between the F3 lobe within the FERM domain of human Myo10 (PDB entry 3AU5) and the cytoplasmic tail of human β1-integrin, align with a RMSD of ∼1.2 Å (see Methods and Fig. S8 for details), (ii) homologous hydrophobic residues in the C-terminal portion of talin’s F3 lobe (mouse talin 2 residues W362, I395, I399, I402, and L403) and Myo10’s F3 lobe (human Myo10 residues F2002, M2033, I2037, I2040, and V2041) make extensive interactions with the NPXY motif in the cytoplasmic tail of β1-integrin [81], (iii) changing Ile 396 in the F3 lobe of chicken talin (and only Ile 396) to an Ala dramatically reduces talin’s affinity for β1-integrin [82], (iv) Ile 396 in chicken talin corresponds to Ile 399 in mouse talin 2, Ile 2037 in human Myo10, and Ile 2041 in mouse Myo10 (the residue we mutated here to a Gln; see Fig. S8 for additional details). mScarlet tagged mut-MyTH4 Myo10 was created by changing four closely-spaced lysine residues present within the mouse Myo10 MyTH4 domain to glutamates (K1651, K1654, K1658, K1661). This positively charged patch has been shown to bind to the acidic tails of α- and β-tubulin, and to be responsible for the ability of the MyTH4 domain to sediment with microtubules [23, 41]. Of note, Kwon et al [43] used human Myo10 in which two of these four lysines were changed to aspartates (K1647 and K1650, which correspond to K1651 and K1654 in mouse Myo10) to identify Myo10 functions that require interaction with microtubules. All clones were confirmed by sequencing.

### Lentivirus packaging and cell transduction

We used the second generation lentiviral packaging plasmid psPAX2 (Addgene, #12260) to generate viral supernatants for pLenti-EB1-EGFP (Addgene, #118084), pLVX-FLAG-dTomato-centrin-1 (Addgene,#73332), pLentiPGK DEST H2B-iRFP670 (Addgene, 90237), pLV-RFP-H2B (Addgene, 26001), and pLenti-EGFP-Cre (Addgene, 86805). These plasmids were co-transfected with psPAX2 and pMD2.G (Addgene, #12259) into HEK-293T cells at ∼80% confluency using LipoD293 (SignaGen, SL100668). Viral supernatants were collected 48 hrs post transfection, clarified by centrifugation at 500 x g for 10 min, and concentrated using a Lenti-X concentrator (TakaraBio, #631231). Viral pellets were obtained by centrifugation of the concentrated virus at 1,500 x g for 45 minutes at 4°C. The pellets were resuspended in complete DMEM and stored in aliquots at −80°C. Purified lentivirus was added directly to cells after the optimal amount was determined by a pilot experiment involving serial dilutions and determining the fraction of cells transduced. After a 6 hrs incubation, the cells were washed 3 times with PBS and returned to complete culture medium for imaging or fixation at various time points. To knock out Myo10 in MEFs isolated from the Myo10 cKO mouse [13], the MEFs were treated with lentivirus expressing EGFP-Cre as described above. Cell phenotypes were determined 48 hrs after lenti-cre transduction. All processes and materials were handled in accordance with the NIH biosafety guidelines.

### Generation of Myo10 KO HeLa cells

To knock out Myo10 in HeLa cells using CRISPR, we used an online CRISPR design tool (http://cripr.mit.edu) to identify the following guide sequences within exon 3 of human Myo10 (NCBI Reference Sequence: NC_000005.10): 5’- CACCGTATGCACCCCACGAACGAGG (PAM)-3’; 3’-CATACGTGGGGTGCTTGCTCCCAAA-5’. These guides sequences were inserted into pSpCas9(BB)-2A-GFP (Addgene, #48138) to create pSpCas9(BB)-2A-GFP-ghMyo10 as described previously [83]. Two days after transfection of WT HeLa cells with pSpCas9(BB)-2A-GFP-ghMyo10 using Amaxa nucleofection (Lonza), GFP-positive cells were subjected to single-cell sorting into 96-well plates using a BD FACS cell sorter. Whole cell protein lysates and genomic DNA prepared from the two single-cell KO clones obtained (KO-1 and KO-2) were characterized by Western blotting and sequencing of PCR products that span the guide sequence sites, respectively.

### Generation of Halo-Myo10 knock-in HeLa cells

The target guide sequence, which is immediately upstream of the starting codon in human Myo10, was designed using an online CRISPR design tool provided by the Zhang lab at MIT (http://cripr.mit.edu):

5’ CACC GGAGCGGCACTCGGCGAGTC (PAM) 3’

3’ CCTCGCCGTGAGCCGCTCAG CAAA 5’

This guide sequence was cloned into pSPCas9(BB)-2A-puro(PX459)V2.0 (Addgene, #62988) as described previously [83] to create pSPCas9(BB)-2A-puro(PX459)-hMyo10. To generate the donor plasmid, a gBlock was designed that contains two HDR (Homology Directed Repair) arms (560 bp 5’ HDR and 460 bp 3’ HDR), the target sequence with the mutation in the PAM site, a mutation to change the initiator methionine in Myo10 to an Alanine, and a multicloning site between the 5’and 3’ HDRs that contains an EcoRI site and an XhoI site. For cloning convenience, ∼20 bp taken from the backbone of plasmid pSP72 (Promega, P2191) was added onto the outside end of both 5’ and 3’HDRs. Below is the complete gBlock sequence (the guide sequence is in bold, the EcoRI and XhoI sites are underlined, and the additional sequences from plasmid pSP72 are in italics).

*ACTGAGAGTGCACCATATG*AGAGCTGGCTGAGCCGCGGCGCGGGACTGCTCACCTCCAAGCGCTCGCG CGGGGATCGCGGCTCCTGCTCACTTTGCGGCCCGCTGTCCTCCTGCCCGCCCCGAGGGCCCCCGGCCGGAGCGCAGAGGGAGGGGGCCGCGCTCGCCAGCACCCCGCCGCCTTCCCCCGCCTGGGGGAAGAATGTGCCACCAGCTGTTCTCCGCTTGCGAGCGCTGCGCCCAGTAGTGAGGAACTTGGAGGAAGAAGAGACAAAGGCTGCCGTCGGGACGGGCGAGTTAGGGACTTGGGTTTGGGCGAACAAAAGGTGAGAAGGACAAGAAGGGACCGGGCGATGGCAGCAGGGGAGCCCCGCGGGCGCGCGTCCTCGGGAGTGGCGCCGTGACACGCA TGGTTTCCCCGGACCCGCGGCGGCGCTGACTTCCGCGAGTC**GGAGCGGCACTCGGCGAGT**CCCGGACT GCGCTGGAACAGCTAGCGCT<u>GAATTC</u>GAGCTGTACAAGTCCGGACTC<u>CTCGAG</u>CCGGATAACTTCTTCA CCGAGGTAAGTGCGCTCCCAGTCCGACCTGGCCTCCGGAGCCCAGGGAGAGAGGCGTCTGCCCACCACGCCGCGCGCCCTGGGTACTTTTTCCTAAGCCCTGGAAGGCGCAACTTTCTGGGAGTCTCCTGAAATCACCCCCCATCCCCCCGCGGAGTCTCTGATGAGTAAGCCCGGGCAGGTTTTGTTTCGTCCTGTCCCGCGCTCGCATTTTGCTCCGGGAGGTAGCGAAGGTGCGTTTCCGTTTGCGTGGGTGGCTGGCTCTCGGGGCGCCCTGG GACACCCGCGCCAGGTGA*AGATCTGCCGGTCTCCCTATA*

This gBlock was cloned into pSP72 using an In-Fusion HD cloning protocol (TakaraBio) to create plasmid pSP72-hMyo10-KI. The Halo tag sequence starting with the Methionine was then inserted into pSP72-hMyo10-KI using the EcoRI and XhoI restriction sites to create the final donor construct, pSP72-Halo-hMyo10-KI. This plasmid and plasmid pSPCas9(BB)-2A-puro(PX459)-hMyo10 above were transfected together into HeLa cells using an Amaxa nucleofection apparatus (Lonza). Five days later the cells were subjected to single-cell sorting into 96-well plates. Several individual clones were analyzed by Western blotting and imaging following incubation for one hour with the cell permeable JF-549 or JF-554 Halo dyes (Janelia/HHMI) at a final concentration of 200 nM (see the text for details).

### Immunoblotting

Whole cell protein lysates were collected directly from culture dishes by adding 1X SDS sample buffer, as described previously [84]. Samples were resolved on 4-12% or 6% NuPAGE Bis-Tris gels (ThermoFisher Scientific) and transferred onto nitrocellulose membranes (Bio-Rad) using a semi-dry transfer system (Bio-Rad). Nitrocellulose membranes were blocked in TBST (10 mM Tris, pH 8.0, 150 mM NaCl, and 0.02% Tween 20) supplemented with 5% milk for 2 hrs, incubated with anti-Myo10 primary antibody overnight at 4°C, washed with TBST, and incubated in secondary antibody at RT for 2 hrs. Antibodies were diluted in TBST containing 5% milk. Actin detected using an anti-β-actin antibody was used as a loading control. Proteins were detected using SuperSignal West Pico Plus Chemiluminescent Substrate (ThermoFisher Scientific) and quantitated using an Amersham Imager 600 (GE Healthcare Life Sciences).

### Growth rate measurements

Primary MEFs isolated from WT C57BL/6 mouse embryos, non-exencephalic Myo10 KO mouse embryos (tm1d), and exencephalic Myo10 KO mouse embryos (tm1d) were seeded in individual wells of a 6-well plate at 2.50 × 10^5^ cells per well for the dense condition, 1.25 × 10^5^ cells per well for the moderate condition, and 2.5 × 10^4^ cells per well for the sparse condition. Cells were counted using Olympus Cell Counter model R1 after 3 days of culture.

### Scoring mitotic phenotypes

To score mitotic phenotypes, HeLa cells (WT, Myo10 KO, Myo10 KD, and Myo10 KO cells rescued with pScarlet-Myo10) and MEFs (isolated from WT C57BL/6 mouse embryos, Myo10 KO mouse embryos (tm1d), and Myo10 cKO mouse embryos (tm1c, scored before and after treatment with pLenti-EGFP-Cre)) were plated in imaging chambers at moderate density, cultured overnight, fixed, and stained with anti-γ-tubulin, anti-α-tubulin and DAPI. Of note, cells were never synchronized for these studies. Z-stack images were taken in 0.25 µm steps. The criteria used to score metaphase cells as bipolar, semi-bipolar, or multipolar are described in the text. The same images were used to quantify the percent of cells exhibiting misaligned chromosomes and lagging chromosomes. Quantitation of the distance in Z between the two spindle poles in cells undergoing bipolar mitosis was based on the number of confocal slices between the γ-tubulin foci marking the two poles. To distinguish mitotic cells containing two centrosomes from mitotic cells containing supernumerary centrosomes, and to distinguish γ-tubulin-positive poles containing two centrioles from those containing one centriole or no centriole, metaphase cells were stained with anti-γ-tubulin, anti-centrin-1 and DAPI. To gauge pole maturation, cells were fixed and stained with anti-CDK5Rap2, anti-centrin-1 and DAPI. Cells with a normal number of centrosomes were imaged in 0.25 µm steps, and the total intensity of CDK5Rap2 per pole was determined by summing slices, then thresholding using the otsu function in Image J.

### Dynamic imaging of acentriolar spindle poles using GFP-EB1

Myo10 KO HeLa cells transduced with pLenti-EB1-EGFP, pLVX-FLAG-dTomato-centrin-1, and pLentiPGK DEST H2B-iRFP670, or pLenti-EB1-EGFP and pLenti-RFP-H2B, were imaged live on an Airyscan 880 microscope equipped with a 60X, 1.4 NA objective (one frame every 2 sec for 2 min).

### Modeling the interaction between the F3 lobe in Myo10’s FERM domain and the cytoplasmic tail of β1-integrin

To predict the interaction between the F3 lobe within the FERM domain of human Myo10 (PDB entry 3AU5) and the cytoplasmic tail of human β1-integrin, we used PyMOL [85] to superimpose the F3 lobe of human Myo10 with the structure of the F3 lobe within the FERM domain of mouse talin 2 FERM domain complexed with the cytoplasmic tail of human β1-integrin (PDB entry 3g9w). F3 lobe residues Y1953 to S2046 in Myo10 and Y311 to S408 in talin 2, and residues G750 to N788 in the β1-integrin cytoplasmic tail, were included in the structural prediction.

### Imaging and statistical analyses

Imaging of both fixed and live cells was performed on a Zeiss Airyscan 880 microscope equipped with a 60X, 1.4 NA objective. Images were processed in auto strength mode using ZenBlack software (Version 2.3) and analyzed using ImageJ. Excel or GraphPad Prism were used for statistical analyses and graphing. Statistical significance was determined using unpaired t-test and indicated as follows: * = P<0.05, ** = P<0.01, *** = P<0.001, and **** = P<0.0001.

## Acknowledgments

This work was supported by the Intramural Research Program of the National Heart, Lung, and Blood Institute (1ZIAHL000514-31 to JAH) and by R01GM134531 to REC. The authors are very grateful to Dr. Mingjie Zhang (HK University of Science and Technology, Hong Kong) for his suggestion to mutate Ile 2041 in mouse Myo10 to attenuate its interaction with integrin. The authors also thank Dr. Nasser Rusan (NHLBI, NIH), Dr. Jadranka Loncarek (NCI, NIH), and Dr. Jeffrey B. Woodruff (UT Southwestern) for their expert advice.

---

**Figure S1. Myo10 protein levels in Myo10 KO HeLa cells, Myo10 KD HeLa cells, and Myo10 cKO MEFs after treatment with lenti-cre.** (A) The amount of Myo10 in KO-1 and KO-2 expressed as a percent of the amount in WT HeLa cells (N= 6; 7.1 ± 5.2% for KO-1 and 13.8 ± 11.5% for KO-2). (B) The amount of Myo10 in WT HeLa cells 12, 24 and 48 hrs after the start of a second round of treatment with Myo10 siRNA, expressed as a percent of the amount in WT HeLa cells treated with a nontargeting siRNA control (N=3; 8.6 ± 2.1% at 12 hr, 3.4 ± 2.1 % at 24 hr, and 0.3 ± 0.1% at 48 hr). (C) The amount of Myo10 in cKO MEFs 6, 12, 24 and 48 hrs after addition of a lentivirus expressing GFP-tagged Cre recombinase, expressed as a percent of the amount in cKO MEFs before the addition of lenti-cre (N=3; 90.3 ± 6.9% at 0 hr, 33.4 ± 8.1% at 6 hr, 24.6 ± 15.5% at 12 hr, 3.9 ± 3.6% at 24 hr, and 3.1 ± 0.7% at 48 hr). In each case, the amount of Myo10 protein was determined by densitometry of Western blots loaded with whole cell extracts and probed with a Myo10 antibody.

**Figure S2. Myo10 KO MEFs exhibit spindle defects whose severity depends upon embryo source and culture density.** (A) Growth rates of WT MEFs, MEFs isolated from non-exencephalic Myo10 KO embryos, and MEFs isolated from exencephalic Myo10 KO embryos and seeded at different cell densities (D, M and S stand for dense, moderate and sparse seeding, respectively; from 3 experiments, each done in duplicate). (B) The same data as in (A) but presented so as to allow comparison of the three types of MEF at each cell density. (C) Quantitation at metaphase of the percent of multipolar spindles in WT MEFs, non-exencephalic Myo10 KO MEFs, and exencephalic Myo10 KO MEFs grown at different cell densities (D: 147 WT MEFs, 145 non-exencephalic KO MEFs, and 186 exencephalic KO MEFs; M: 142 WT MEFs, 177 non-exencephalic KO MEFs, and 241 exencephalic KO MEFs; S: 161 WT MEFs, 209 non-exencephalic KO MEFs, and 257 exencephalic KO MEFs; from 4 experiments). (D) The same data as in (C) but presented so as to allow comparison of the three types of MEF at each cell density. (E) Average distance in Z at metaphase between the two poles of WT MEFs, non-exencephalic Myo10 KO MEFs, and exencephalic Myo10 KO seeded at moderate density. (F) The distributions of the values in (E) in 1 µm intervals.

**Figure S3. Validation of the anti-Myo10 antibody.** (A1-A5) Metaphase WT HeLa cell fixed and stained with anti-Myo10, Phalloidin and DAPI and imaged to show endogenous Myo10 at the tips of retraction fibers. (B1-B5) Same as A1-A5 except a metaphase KO-1 cell. The mag bars are in A4 and B4 are 10 µm. The mag bars in A5 and B5 are 2 µm.

**Figure S4. Halo-Myo10 KI and validation.** (A) Western blot of a WT HeLa cell extract (lane 1) and a Halo-Myo10 KI cell extract (lane 2) probed with anti-Myo10 antibody and anti-GAPDH as a loading control. (B1, B2) An interphase Halo-Myo10 KI cell fixed and stained with Alexa488- labeled Phalloidin and DAPI. (C1-C5) An interphase Halo-Myo10 KI cells fixed and stained with anti-Myo10 and DAPI. The mag bars in B1 and C4 are 10 µm. The mag bars in B2 and C5 are 5 µm.

**Figure S5. Cytokinesis failure and centriole overduplication do not contribute to the multipolar phenotype exhibited by Myo10 KO HeLa cells.** (A1 and A2) Unsynchronized WT HeLa, KO-1 and KO-2 were stained with Phalloidin and DAPI (A1 shows the staining of KO-1) and the number of nuclei per cell quantified (A2) (315 WT HeLa, 313 KO-1 cells, and 307 KO-2 cells from 3 experiments; 1.1 ± 0.3 nuclei per cell for WT, 1.0 ± 0.2 nuclei per cell for KO-1, and 1.1 ± 0.3 nuclei per cell for KO-2). (B) Unsynchronized WT HeLa, KO-1 and KO-2 were stained with centrin-1 and the number of centrioles per cell quantified (155 WT HeLa, 154 KO-1 cells, and 172 KO-2 cells from 3 experiments; 3.1 ± 1.6 centrioles per cell for WT, 2.9 ± 1.2 centrioles per cell for KO-1, and 2.9 ± 1.1 centrioles per cell for KO-2). The mag bar is 40 µm.

**Figure S6. Extreme examples Myo10 KO HeLa cells possessing multiple, presumptive, centriole-negative spindle poles generated by PCM fragmentation.** (A and B) Two examples of KO-1 cells stained for γ-tubulin and centrin-1 that exhibit many presumptive, centriole-negative spindle poles generated by PCM fragmentation in addition to two (A) or four (B) normal poles containing two centriole each (see also Z-Stack Movie 18 for (A) and Z-Stack Movie 19 for (B)). All the mag bars are 1 µm.

**Figure S7. Like WT Myo10, mut-FERM Myo10 and mut-MyTH4 Myo10 localize at the tips of interphase filopodia and metaphase retraction fibers.** Representative examples of interphase KO-1 cells rescued with mScarlet-WT Myo10 (A1-A3), mScarlet-mut-FERM Myo10 (B1-B3) or mScarlet-mut-MyTH4 Myo10 (C1-C3) that were fixed and stained with phalloidin. Representative examples of metaphase KO-1 cells rescued with mScarlet-WT Myo10 (D1-D4), mScarlet-mut-FERM Myo10 (E1-E4) or mScarlet-mut-MyTH4 Myo10 (F1-F4) that were fixed and stained with phalloidin. Magnified insets are shown for all of the images. The mag bars in A3, B3 and C3 are 10 µm. The mag bars in D4, E4 and F4 are 5 µm.

**Figure S8. Predicted interaction between the F3 lobe within Myo10’s FERM domain and the cytoplasmic tail of β1-integrin** (A) The structure of the F3 lobe within the FERM domain of mouse talin 2 complexed with the cytoplasmic tail of human β1-integrin (PDB entry 3G9W). (B) Predicted structure of the complex between the F3 lobe with the FERM domain of human Myo10 (PDB entry 3AU5) and the cytoplasmic tail of human β1-integrin (see Methods for details). The hydrophobic residues within the F3 lobes of talin 2 and Myo10 that drive interaction with the NPXY motif in β1-integrin [81] are noted in green and blue, respectively. The β1-integrin cytoplasmic tail is colored dark violet. The two structures align with a RMSD of ∼1.2 Å. (C) Shown is an alignment between the hydrophobic residues within the C-terminal portion of the F3 lobes of chicken talin, mouse talin 2, human Myo10, and mouse Myo10 that drive interaction with the cytoplasmic tail of β1-integrin. The hydrophobic residues within the F3 lobes of mouse talin 2 and human Myo10 that are noted in A and B are underlined. The isoleucine in chicken talin 2 (Ile 396), whose mutation to an alanine residue dramatically decreases talin 2’s affinity for β1-integrin [82], and the corresponding isoleucine that we mutated to a glutamine residue in mouse Myo10 (Ile 2041) to create mut-FERM Myo10, are marked with asterisks.

## Movie Legends

**Movie 1**. Z-Stack movie of a bipolar cell at metaphase that was stained with anti-γ-tubulin-Cy3, anti-tubulin-AF488 and DAPI, and sectioned in 0.25 µm intervals. Played at 5 fps. The mag bar is 5 µm.

**Movie 2**. Z-Stack movie of a semi-bipolar cell at metaphase that was stained with anti-γ- tubulin-Cy3, anti-tubulin-AF488 and DAPI, and sectioned in 0.25 µm intervals. Played at 5 fps. The mag bar is 5 µm.

**Movie 3**. Z-Stack movie of a multipolar cell at metaphase that was stained with anti-γ-tubulin-Cy3, anti-tubulin-AF488 and DAPI, and sectioned in 0.25 µm intervals. Played at 5 fps. The mag bar is 5 µm.

**Movie 4**. Z-Stack movie of a bipolar Myo10 KO-1 cell at metaphase that was stained with anti-γ- tubulin-Cy3, anti-tubulin-AF488 and DAPI, and sectioned in 0.25 µm intervals. Note that the cell’s two spindle poles are in different Z-planes. Played at 5 fps. The mag bar is 5 µm.

**Movie 5**. Z-Stack movie of a bipolar Myo10 KO-1 cell at anaphase that was stained with ant-γ- tubulin-Cy3, anti-tubulin-AF488 and DAPI, and sectioned in 0.25 µm intervals. Note that the cell’s two spindle poles are in different Z-planes. Played at 5 fps. The mag bar is 5 µm.

**Movie 6**. Z-Stack movie of a bipolar Myo10 KO-1 cell at telophase that was stained with anti-γ- tubulin-Cy3,anti-tubulin-AF488 and DAPI, and sectioned in 0.25 µm intervals. Note that the cell’s two spindle poles are in different Z-planes. Played at 5 fps. The mag bar is 5 µm.

**Movie 7**. Z-Stack movie of a metaphase Myo10 KO-1 cell expressing mScarlet-Myo10 that was fixed and stained with Alexa488-labeled Phalloidin and DAPI, and sectioned in 0.25 µm intervals to show that mScarlet-Myo10 localizes at the tips of retraction fibers and dorsal filopodia, but not at spindle poles or in the equatorial cortex. Played at 10 fps. The mag bar is 5 µm.

**Movie 8**. Z-Stack movie of a metaphase Myo10 KO-1 cell expressing mScarlet-Myo10 that was fixed and stained with anti-γ-tubulin-AF488 and DAPI, and sectioned in 0.25 µm intervals to show that mScarlet-Myo10 is not enriched at either spindle poles or the equatorial cortex. Played at 10 fps. The mag bar is 5 µm.

**Movie 9**. Z-Stack movie of a metaphase cell that was stained with anti-Myo10-Cy3, Alexa488- labeled Phalloidin and DAPI, and sectioned in 0.25 µm intervals to show that endogenous Myo10 localizes at the tips of retraction fibers and dorsal filopodia, but not at spindle poles or in the equatorial cortex. Played at 10 fps. The mag bar is 5 µm.

**Movie 10.** Z-Stack movie of a metaphase cell that was stained with anti-Myo10-Cy3, anti-γ- tubulin-AF488 and DAPI, and sectioned in 0.25 µm intervals to show that endogenous Myo10 is not enriched at either spindle poles or the equatorial cortex. Played at 10 fps. The mag bar is 5 µm.

**Movie 11.** Z-Stack movie of a metaphase Halo-Myo10 KI cell that was labeled with the JF-SF554 Halo dye, stained with Alexa488-labeled Phalloidin and DAPI, and sectioned in 0.25 µm intervals to show that endogenously tagged Myo10 localizes at the tips of retraction fibers and dorsal filopodia, but not at spindle poles or in the equatorial cortex. Played at 10 fps. The mag bar is 5 µm.

**Movie 12.** Z-Stack movie of a metaphase Halo-Myo10 KI cell that was labeled with the JF-SF554 Halo dye, stained with anti-γ-tubulin-AF488 and DAPI, and sectioned in 0.25 µm intervals to show that endogenously tagged Myo10 is not enriched at either spindle poles or the equatorial cortex. Played at 10 fps. The mag bar is 5 µm.

**Movie 13.** Time lapse movie of a dividing, JF-SF554 Halo dye-labeled Halo-Myo10 KI cell expressing the F-actin reporter GFP-F-Tractin. Shown is an equatorial plane of this cell from metaphase to anaphase. Images were taken once every 60 seconds for 40 minutes and played back at 10 fps. Note that endogenously tagged Myo10 is not enriched at the equatorial cortex during either metaphase or anaphase. The mag bar is 5 µm.

**Movie 14.** Z-Stack movie of a metaphase, JF-SF554 Halo dye-labeled Halo-Myo10 KI cell that had been plated on an I bar pattern of fibronectin, stained with Alexa488-labeled Phalloidin and DAPI, and sectioned in 0.25 µm intervals. Note that endogenously tagged Myo10 is not enriched within that portion of the equatorial cortex where the retraction fibers are concentrated. Played at 10 fps. The mag bar is 5 µm.

**Movie 15.** Z-Stack movie of a metaphase Myo10 KO-1 cell expressing mScarlet-Myo10 that had been plated on an I bar pattern of fibronectin, stained with Alexa488-labeled Phalloidin and DAPI, and sectioned in 0.25 µm intervals. Note that mScarlet-Myo10 is not enriched within that portion of the equatorial cortex where the retraction fibers are concentrated. Played at 10 fps. The mag bar is 5 µm.

**Movie 16.** Z-Stack movie of a Myo10 KO-1 cell exhibiting a multipolar spindle produced by centriolar disengagement alone that was stained with anti-γ-tubulin-Cy3, anti-centrin-1-AF488 and DAPI, and sectioned in 0.25 µm intervals. Played at 5 fps. The mag bar is 5 µm.

**Movie 17.** Z-Stack movie of a Myo10 KO-1 cell exhibiting a multipolar spindle produced by PCM fragmentation alone that was stained with anti-γ-tubulin-Cy3, anti-centrin-1-AF488 and DAPI, and sectioned in 0.25 µm intervals. Played at 5 fps. The mag bar is 5 µm.

**Movie 18.** Z-Stack movie of a multipolar Myo10 KO-1 cell with acentriolar poles generated by PCM fragmentation alone that was stained with anti-γ-tubulin-Cy3, anti-centrin-1-AF488 and DAPI, and sectioned in 0.25 µm intervals. Played at 5 fps. The mag bar is 5 µm.

**Movie 19.** Z-Stack movie of a multipolar Myo10 KO-1 cell with acentriolar poles generated by PCM fragmentation along with centrosome de-clustering that was stained with anti-γ-tubulin-Cy3, anti-centrin-1-AF488 and DAPI, and sectioned in 0.25 µm intervals. Played at 5 fps. The mag bar is 5 µm.

**Movie 20.** Live Z-Stack movie of a bipolar Myo10 KO-1 cell with two centriolar poles (2 centrioles/pole) that was expressing EB1-EGFP, dTomato-centrin-1 and H2B-iRFP670, and sectioned in 0.25 µm intervals. Played at 5 fps. The mag bar is 5 µm.

**Movie 21.** Live Z-Stack movie of a multipolar Myo10 KO-1 cell with three centriolar poles (2 centrioles/pole) that was expressing EB1-EGFP, dTomato-centrin-1 and H2B-iRFP670, and sectioned in 0.25 µm intervals. Played at 5 fps. The mag bar is 5 µm.

**Movie 22.** Live Z-Stack movie of a multipolar Myo10 KO-1 cell with three centriolar poles (2 centrioles/pole) and three acentriolar poles that was expressing EB1-EGFP, dTomato-centrin-1 and H2B-iRFP670, and sectioned in 0.25 µm intervals. Played at 5 fps. The mag bar is 5 µm.

**Movie 23.** Time-lapse images of EB1-EGFP at a centriolar pole in the Myo10 KO-1 cell shown in Movie 22. Images were taken every 2 seconds for 2 minutes and are played back at 5 fps. The mag bar is 5 µm.

**Movie 24.** Time-lapse images of EB1-EGFP at an acentriolar pole in the Myo10 KO-1 cell shown in Movie 22. Images were taken every 2 seconds for 2 minutes and are played back at 5 fps. The mag bar is 5 µm.

**Movie 25.** Time lapse images of a Myo10 KO-1 cell expressing EB1-EGFP and H2B-iRFP670 that is exhibiting PCM/pole fragmentation at metaphase. To create this movie, the equatorial portion of the cell was optically sectioned in 4, 0.5 µm steps every 3 seconds for 60 minutes. The 4 contiguous sections at each time point (spanning 2 µm) were then stacked by maximum intensity projection and played back at 500 fps. Note that the newly-generated EB1-positive pole moves away from the major pole that spawned it, and that it then creates a distortion in the alignment of chromosomes at the metaphase plate as the cell progresses into anaphase. The mag bar is 5 µm.

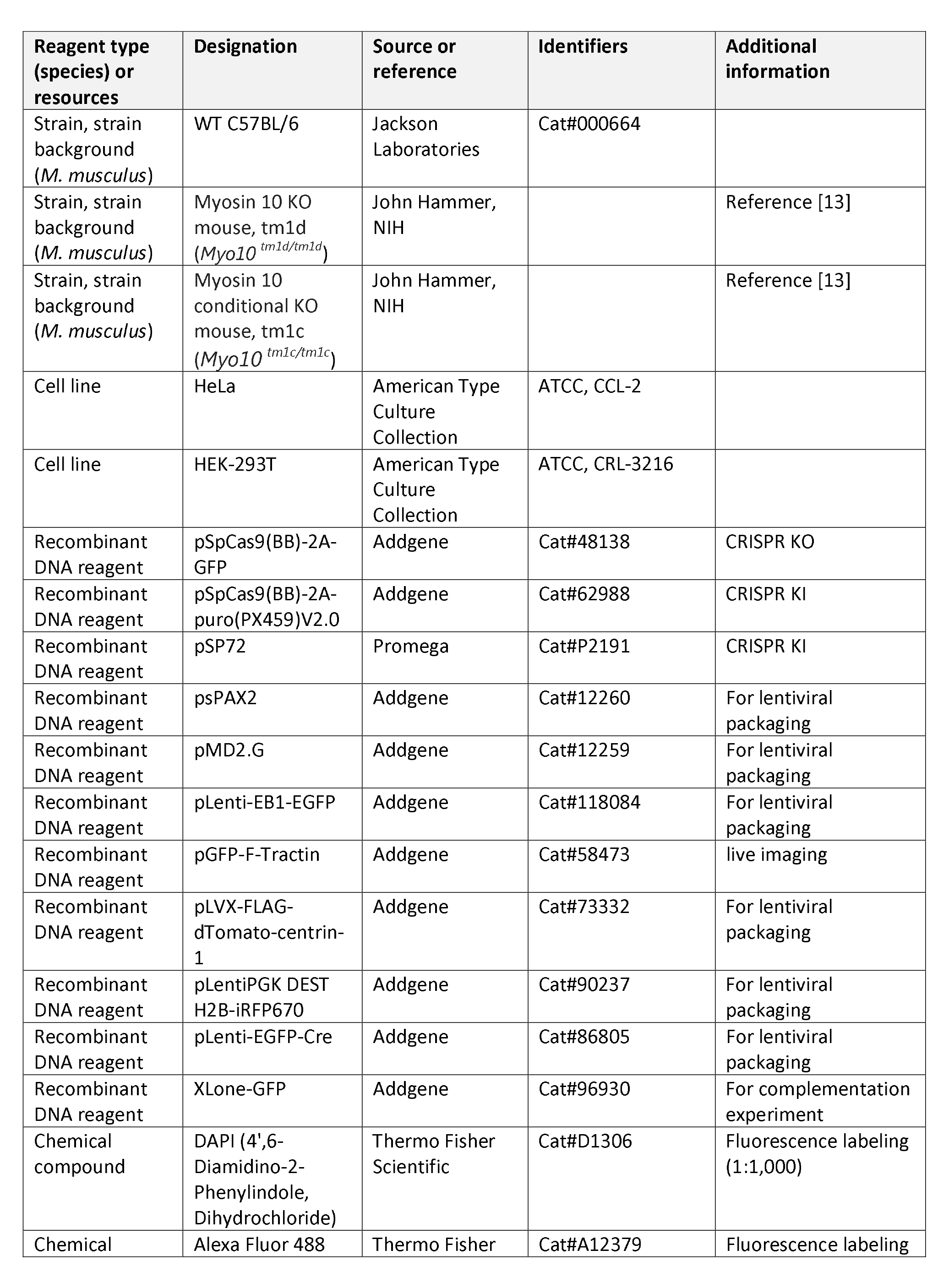
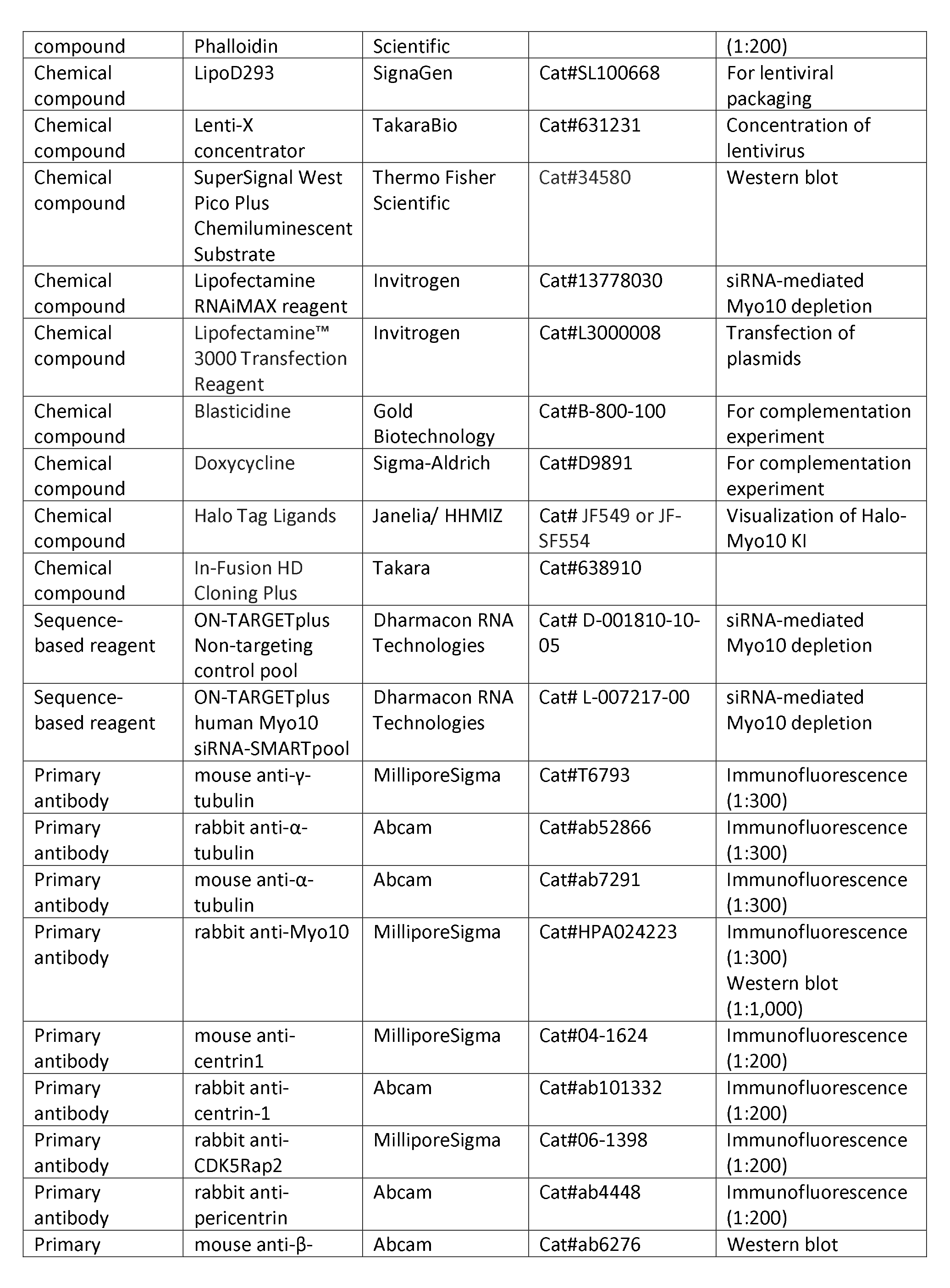
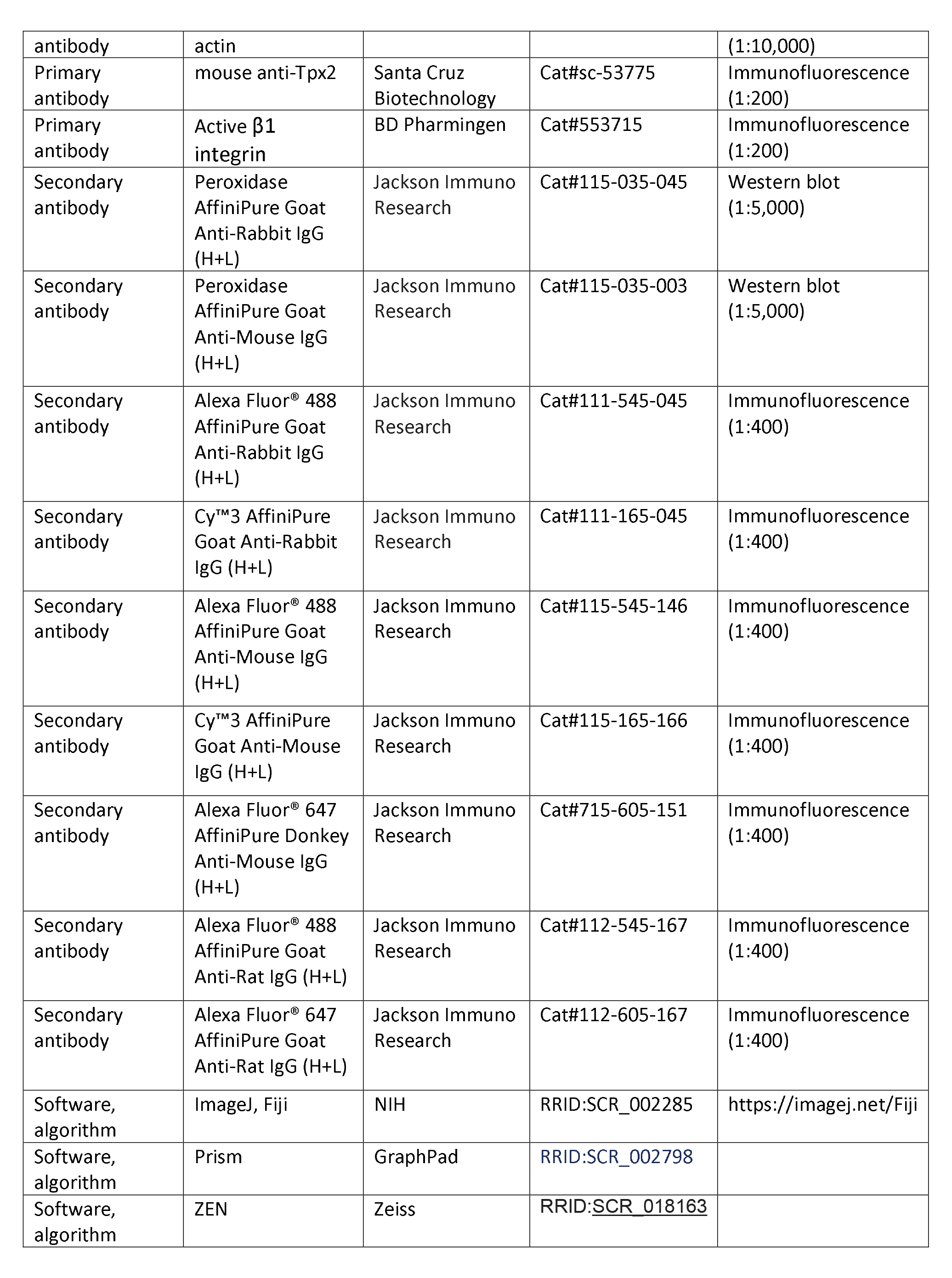

## Notes

### Competing Interest Statement

The authors have declared no competing interest.

### Summary of Updates

uploading all the movies missing in the previous submission

