## supplemental figures for "Myosin 10 uses its MyTH4 and FERM domains differentially to support two aspects of spindle pole biology required for mitotic spindle bipolarity"

FigS1

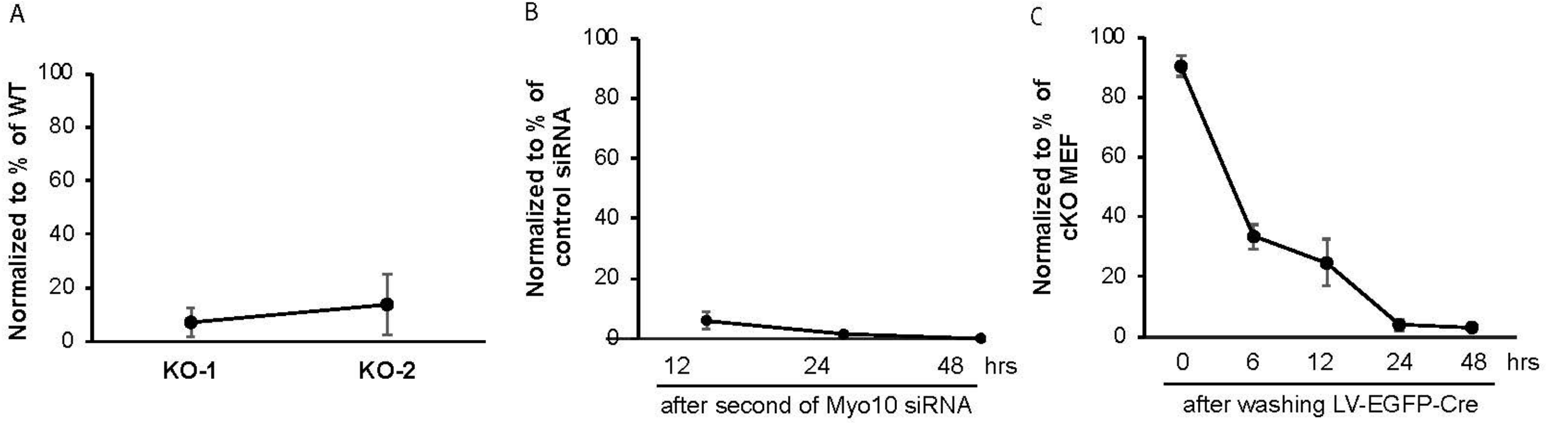
A
100
80
60
40
20
0
Normalized to % of WT
KO-1
KO-2
B
100
80
60
40
20
0
Normalized to % of control siRNA
12
24
48
hrs
after second of Myo10 siRNA
C
100
80
60
40
20
0
Normalized to % of cKO MEF
0
6
12
24
48
hrs
after washing LV-EGFP-Cre

FigS2

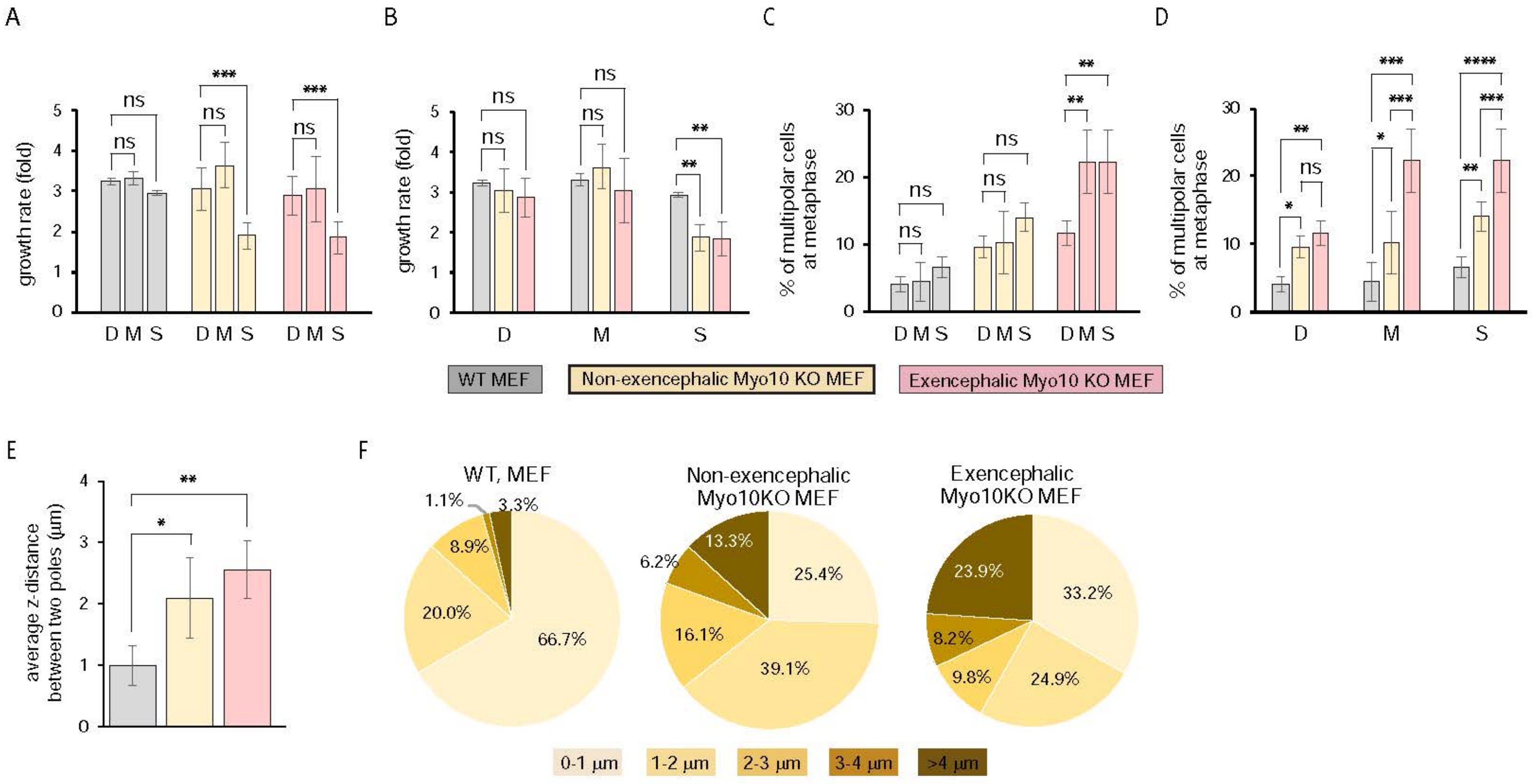
A
B
C
D
growth rate (fold)
% of multipolar cells at metaphase
ns
***
**
****
*
D M S
D
M
S
WT MEF
Non-exencephalic Myo10 KO MEF
Exencephalic Myo10 KO MEF
E
F
average z-distance between two poles (μm)
WT, MEF
Non-exencephalic Myo10KO MEF
Exencephalic Myo10KO MEF
1.1%
3.3%
8.9%
20.0%
66.7%
13.3%
6.2%
25.4%
16.1%
39.1%
23.9%
33.2%
8.2%
9.8%
24.9%
0-1 μm
1-2 μm
2-3 μm
3-4 μm
>4 μm

FigS3

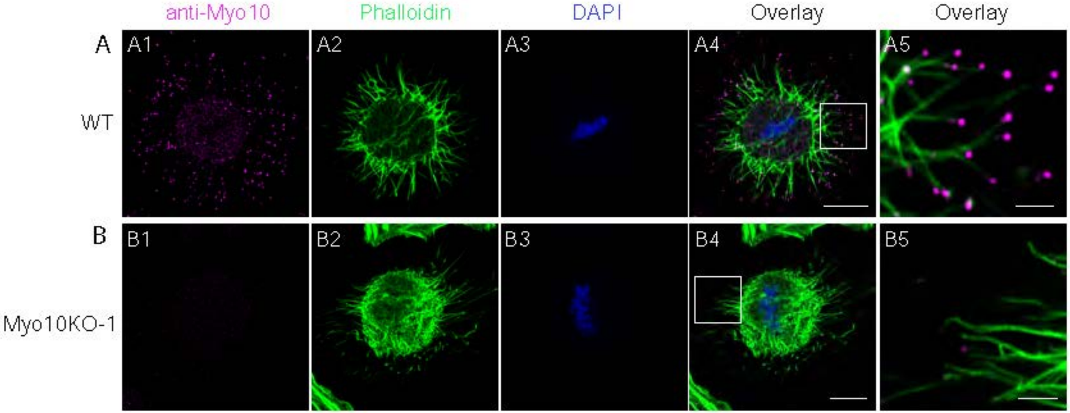
anti-Myo10
Phalloidin
DAPI
Overlay
Overlay
A
WT
A1
A2
A3
A4
A5
B
Myo10KO-1
B1
B2
B3
B4
B5

FigS4

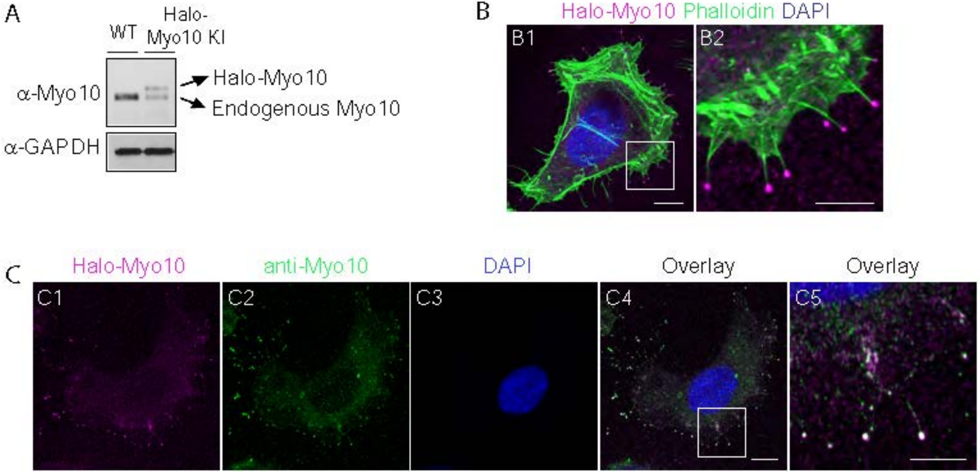
A
WT
Halo-Myo10 KI
α-Myo10
α-GAPDH
Halo-Myo10
Endogenous Myo10
B
Halo-Myo10 Phalloidin DAPI
B1
B2
C
Halo-Myo10
anti-Myo10
DAPI
Overlay
Overlay
C1
C2
C3
C4
C5

### FigS5

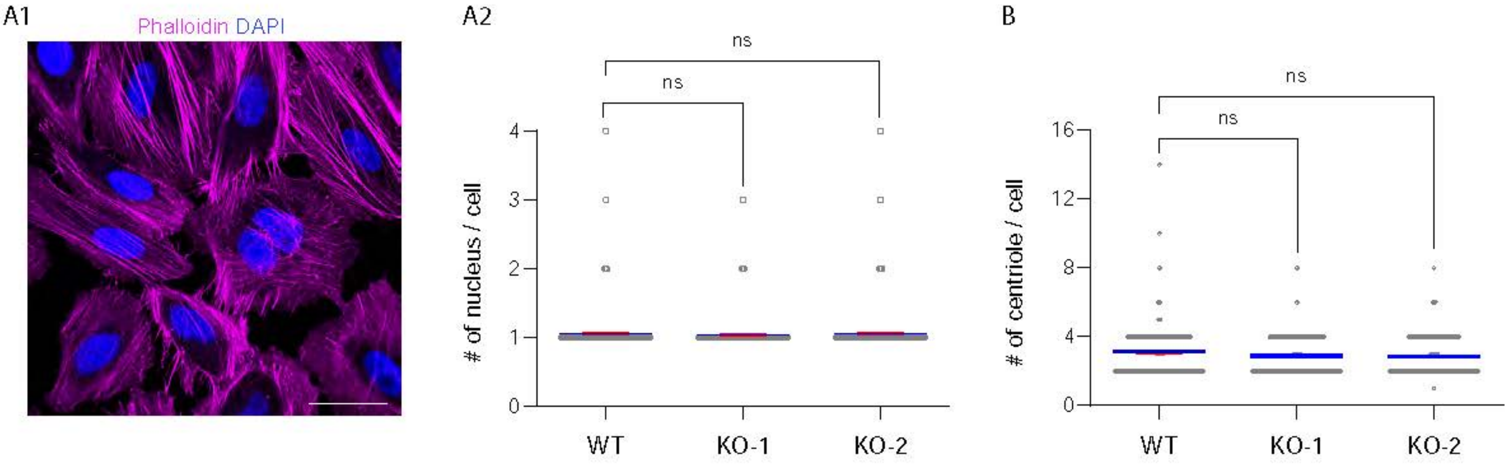
A1
Phalloidin DAPI
A2
ns
ns
### of nucleus / cell
0
1
2
3
4
WT
KO-1
KO-2
B
ns
ns
### of centriole / cell
0
4
8
12
16
WT
KO-1
KO-2

FigS6

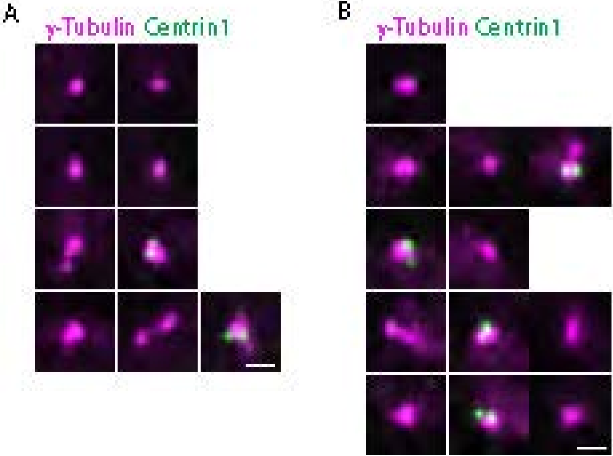

FigS7

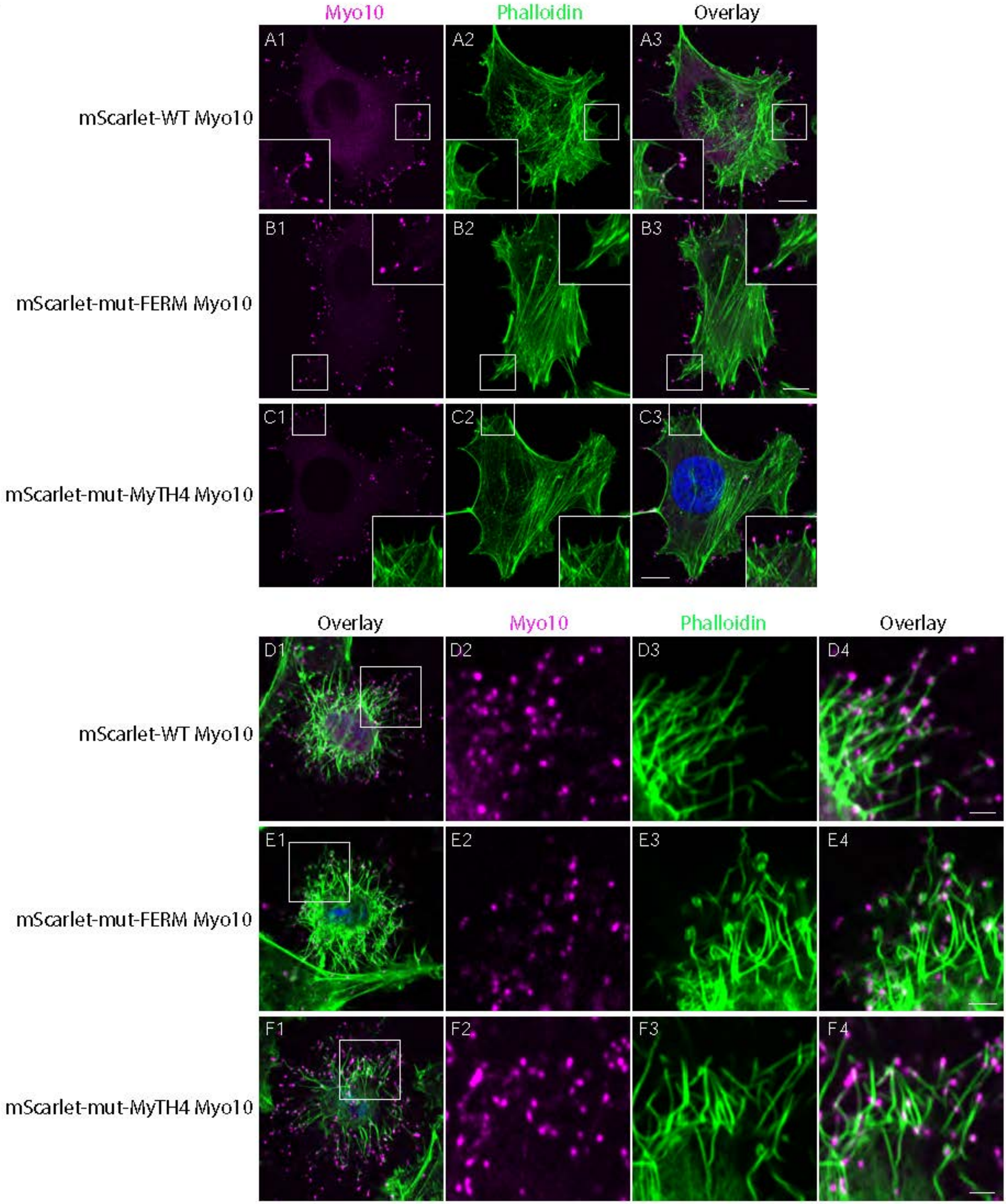
Myo10
Phalloidin
Overlay
A1
A2
A3
mScarlet-WT Myo10
B1
B2
B3
mScarlet-mut-FERM Myo10
C1
C2
C3
mScarlet-mut-MyTH4 Myo10
Overlay
Myo10
Phalloidin
Overlay
D1
D2
D3
D4
mScarlet-WT Myo10
E1
E2
E3
E4
mScarlet-mut-FERM Myo10
F1
F2
F3
F4
mScarlet-mut-MyTH4 Myo10

FigS8

A Talin-2 FERM and Integrin

B Myosin-X FERM and Integrin

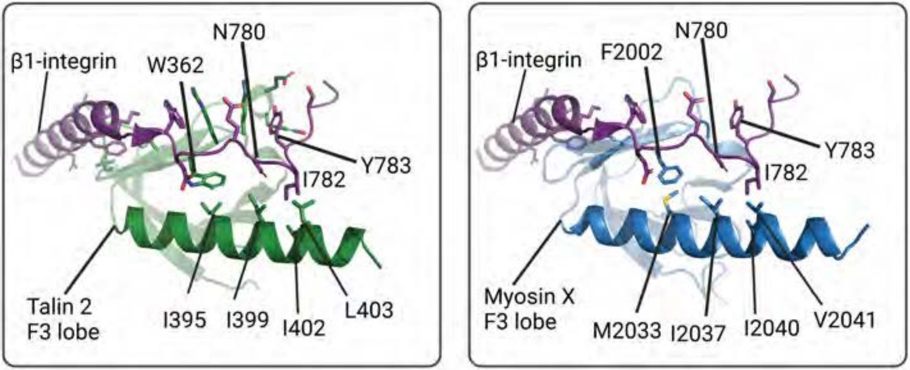


C

I396A

| | | | | | | | | | | | |
|---|---|---|---|---|---|---|---|---|---|---|---|
| | | | | | | * | | | | | |
| cTalin | 392 | I | A | G | Y | I | D | I | I | L | 400 |
| mTalin2 | 395 | I | A | G | Y | I | D | I | I | L | 403 |
| hMyo10 | 2033 | M | K | A | Y | I | S | M | I | V | 2041 |
| mMyo10 | 2037 | M | K | A | Y | I | S | M | I | V | 2045 |
| | | | | | | * | | | | | |

I2041Q
